# IspE Kinase as an Anti-infective Target: Role of a Hydrophobic Pocket in Inhibitor Binding

**DOI:** 10.1101/2024.06.17.599194

**Authors:** Rawia Hamid, Danica J. Walsh, Eleonora Diamanti, Diana Aguilar, Antoine Lacour, Mostafa M. Hamed, Anna K. H. Hirsch

## Abstract

Enzymes of the methylerythritol phosphate (MEP) pathway are potential targets for antimicrobial drug discovery. Here we focus on 4-diphosphocytidyl-2-C-methyl-D-erythritol (IspE) kinase from the MEP pathway. We use biochemical and structural biology methods to investigate homologs from the pathogenic microorganisms; *Escherichia coli, Klebsiella pneumoniae*, and *Acinetobacter baumannii*. We determined the X-ray structures of IspE-inhibitor complexes and studied selected inhibitors’ binding modes targeting the substrate pocket. The experimental results indicate the need for distinct inhibitor strategies due to structural differences among IspE homologs, particularly for A. baumannii IspE, which displays unique inhibitory profile due to a tighter hydrophobic subpocket in the substrate binding site. This study enhances our understanding of the MEP enzymes and sets the stage for structure-based drug design of selective inhibitors to combat pathogenic microorganisms.

## 1 INTRODUCTION

There is a growing need for new anti-infectives. The world is racing to identify new microbial targets to combat multidrug–resistant pathogens ^1,2^. The 2*C*-Methyl-D-erythritol 4-phosphate (MEP) pathway is a rich source of attractive underexplored anti-infective targets ^3,4^ due to its role in the production of the universal precursors of isoprenoids, especially in pathogenic microorganisms ^5,6^. Unlike humans, many of these microorganisms rely on the MEP pathway for the synthesis of the isoprenoids building blocks; isopentenyl diphosphate (IDP) and dimethylallyl diphosphate (DMADP). Several Gram-negative bacteria, like *Escherichia coli, Klebsiella pneumoniae and Acinetobacter baumannii* depend on this pathway for isoprenoid production ^7,8^. Constituent enzymes of the MEP pathway are validated anti-infective drug targets as confirmed through the *in-vivo* efficacy of the antimalarial drug fosmidomycin, which targets the pathway’s second step catalyzed by the enzyme IspC ^9,10^.

This study focuses on the only kinase present in the MEP pathway, the so-called IspE. The IspE gene (ychB) was first identified in 1999 by Lüttgen et al. as the enzyme that follows IspD in the MEP pathway. They provided evidence of ATP-dependent phosphorylation of the substrate 4-diphosphocytidyl-2-*C*-methyl-D-erythritol (CDP-ME) ^11^ (Fig.1). Building upon these findings, further investigations led by Kuzuyama and his team, confirmed the role of the ychB gene and provided additional insights into its kinetic product ^12^. Until now, IspE crystal structures have been elucidated for *Aquifelix aeolicus* IspE (*Aa*IspE, PDB ID: 2VF3) ^13^, *E. coli* IspE (*EcIspE*, PDB ID: 10J4) ^14^, *Thermus thermophilus* (*Tt*IspE, PDB Id: 1UEK) ^15^ and *Mycobacterium tuberculosis* (*Mtb*IspE, PDB ID: 3PYD) ^16^.

**Figure 1:**
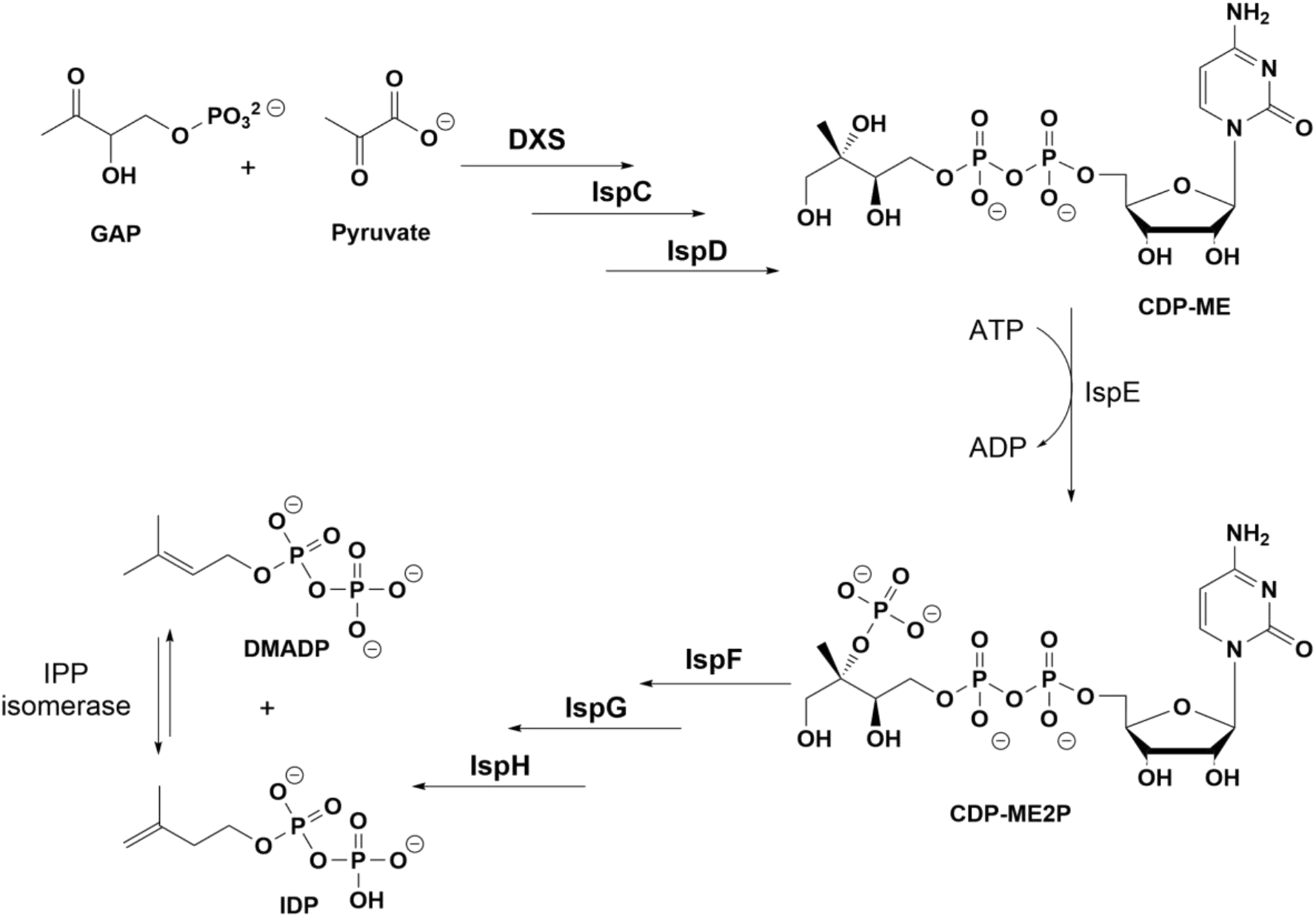
MEP pathway, highlighting the reaction catalyzed by IspE kinase.

IspE kinase, an ATP-dependent enzyme, shares structural similarities with the galactokinase, homoserine kinase, mevalonate kinase and phosphomevalonate kinase (GHMP) enzymes. The active site of this family is characterized by five regions responsible for binding the substrate CDP-ME and the co-factor ATP. These regions interact specifically with adenosine, methyl-erythritol, cytidine, and phosphate groups. Additionally, there is a small hydrophobic pocket that is partially accommodated by the methyl group of the substrate CDP-ME ^13–15^. In contrast to most GHMP kinases IspE is mostly present in solution as a monomer ^17^. Two available structures of *E. coli* IspE show two molecules in the asymmetric unit (ASU). Notably, each crystal lattice configuration is distinct. A structure of IspE in complex with CDP-ME and ATP (PDB-ID:1OJ4) has a 2-fold symmetry ^14^. On the other hand, a complex structure of IspE and ADP (PDB ID: 2WW4), lacks this symmetry and parts of both molecules in the ASU are found too close to each other’s active site, specifically the CDP-ME pocket ^18^. This proximity leads to enzyme inactivity *in vivo* because of unavailable substrate pocket. This suggests that the formation of two molecules in the ASU is an outcome of the crystallization process itself. This arrangement seems to be predominant in the IspE crystal lattice, which poses a challenge in drug design. Attempting to co-crystallize/soak inhibitors targeting the substrate-binding site to elucidate their binding mode is hampered by an essentially unavailable pocket.

Here we focus on the structural analysis of IspE from human pathogens, namely, *Escherichia coli, Klebsiella pneumoniae*, and *Acinetobacter baumannii*. We study the interactions between the ligand and the active-site of IspE from these three pathogens. By investigating these aspects, the study seeks to gain a deeper understanding of the IspE enzymes in these human pathogens and their potential as targets for developing new therapeutic interventions.

## 2 RESULTS AND DISCUSSION

### 2.1 Kinetic Characterization of IspE from *Escherichia coli, Klebsiella pneumoniae* and *Acinetobacter baumannii*

To characterize IspE kinase homologs with respect to their inhibitor binding properties, the plasmid containing IspE genes for expressing IspE from *E. coli, K. pneumoniae*, and *A. baumannii* (from BioCat GmbH) were transformed into *E. coli* B21(DE3), the corresponding proteins overexpressed, purified and their quality analyzed. The activities of the isolated enzymes from the three pathogens were measured by coupling the IspE kinase activity to the auxiliary enzymes lactate dehydrogenase and pyruvate kinase, using a spectrophotometric assay that quantifies the reduction in the absorption of NADH at 340 nm ^19^.

*E. coli* IspE showed a *K*_m_^CDP-ME^ of 200 μM (V_max_ 171 µM min^−1^) and *K*_m_^ATP^ of 420 μM (V_max_ 195 µM min^−1^). For *K. pneumoniae* IspE the apparent *K*_m_^CDP-ME^ was 170 µM (V_max_ 119 µM min^−1^) and *K*_m_^ATP^ is 348 µM (V_max_ 121 µM min^−1^). These kinetic values are in the same range as those reported previously ^16,19,20^. *E. coli* and *K. pneumoniae* IspE share a sequence homology of 84%, which is also reflected in their kinetic and structural similarity (discussed below). *A. baumannii* IspE shows comparable values with a *K*_*m*_^ATP^of 474 µM (V_max_ 153 µM min^−1^), however, it shows a slightly higher values of *K*_*m*_^CDP-ME^: 358 µM (V_max_ 112 µM min^−1^) (Fig. S1). Noteworthy, *A. baumannii* IspE shares only 39% sequence identity with *E. coli* and 42% with *K. pneumoniae*. Specifically, the sequence alignment from the three homologs reported in Fig. S2 highlights the amino acids similarities and differences in the active site that might have contributed to the different patterns of ligand affinities between these homologs (also discussed below). Most prominent are the change of Pro182 in *E. coli* IspE and *K. pneumoniae* IspE into Gln174 in *A. baumannii* IspE, and the change of Cys211 to Phe205 in *A. baumannii* IspE. These two residues are important for binding to the ribose moiety and methyl of the substrate CDP–ME, respectively.

Gel-filtration analysis shows that all three homologs exist mainly as monomers in solution, with a small amount in an oligomeric state (Fig. S3). This observation agrees with earlier studies on other IspE kinase homologs and indicate that IspE operates as a monomer, with the active site formed within a single polypeptide unit, without any evident involvement from another subunit in the catalytic process ^13,18,21^. The determined biochemical and kinetic values were subsequently used to optimize assay conditions for each homolog to evaluate inhibitory activity.

### 2.1 Molecular structure of *K. pneumoniae* IspE is highly similar to *E. coli* IspE, but crystal packing conceals substrate binding pocket

The 3D structure of *K. pneumoniae IspE* was determined at a resolution of 1.8 Å using molecular replacement and further refined to R-work and R-free values of 19% and 22%, respectively (PDB entry: 8CKH). *E. coli* and *K. pneumoniae* both belong to the Enterobacteriaceae family, with an 84% sequence identity. Predictably, they show similar structural features reflected in an RMSD of 0.557 Å. In addition, conserved residues between the two homologs are involved in binding to ATP and the substrate CDP-ME.

The orthorhombic *K. pneumoniae* IspE crystals contain two molecules in the asymmetric unit that bury an interface of 1167 Å^2^ by a total solvent accessible surface area of 24,927 Å^2^. Strands β2 and β3 and the connecting turn from one molecule point into the substrate binding site of another molecule. This site is occupied by the phosphate of the substrate CDP-ME in the monoclinic crystal form of *E. coli* IspE. The association between the two molecules in *K. pneumoniae* IspE seems to be promoted by a hydrogen bond between the side chain of Arg22 and the main chain of Leu137, with an average distance of 2.8 Å. Such packing of the two molecules prevents substrate or inhibitor binding (Fig. 2).

**Figure 2:**
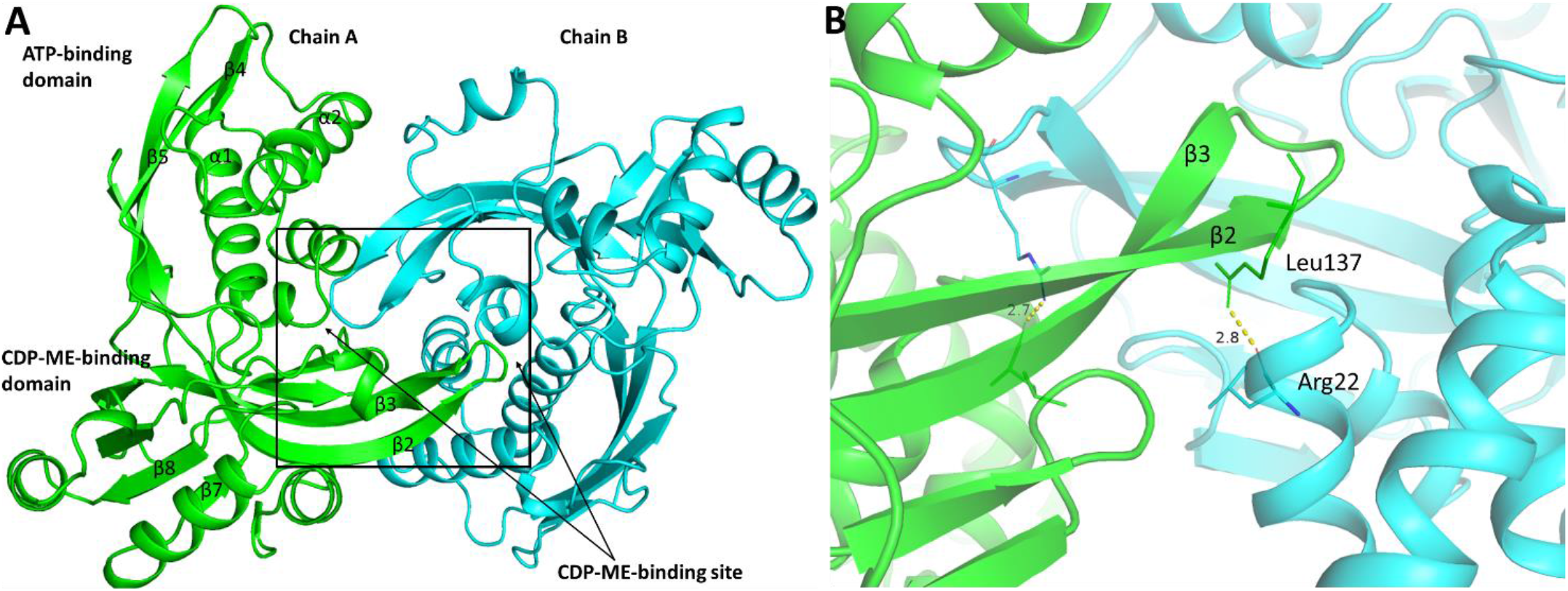
Structure of *Klebsiella pneumoniae* IspE: subunits and domains organization with molecule A shown in green and molecule B in cyan. **A:** domain organization: *N*-terminal, ATP-binding domain has four strands and three helices. The ATP binding site is bordering helices 1 and 2. The CDP-ME binding domain is made up of eight strands and five helices in the C-terminal domain. β2 and β3 in the C-terminal domain are antiparallel to each other and they are curved to form the main part of the substrate binding side. **B:** close up visualization of the clash the two subunits inflicted on each other where the β2-β3 turn is sliding in the CDP-ME binding pocket. Hydrogen bonds are formed between Arg22 side chain of one molecule and main chain in Leu137 of the other molecule (distance 2.8 Å). Interacting residues are shown as sticks: C: green and cyan; O: red; N: blue; S: yellow.

Similar crystallization artifacts have been observed before in the triclinic form of *E. coli* IspE ^18^. This observation holds particular significance in drug design, especially when attempting to co-crystallize inhibitors targeting the substrate-binding pocket. In this study, for the first time, we experimentally show that the structural and kinetic values of *K. pneumoniae* IspE prove its high similarity to *E. coli* IspE, suggesting that these two enzymes share a similar way of ligand binding. These findings led to the assumption that they can be used interchangeably for drug-discovery projects, with the recommendation to prioritize the more robust homolog.

### 2.2 Monoclinic structure of *E. coli* IspE: a novel crystal form exhibiting a distinct arrangement of the two subunits that allows ligand binding

The new crystals of *E. coli* IspE found by us align with the overall structure identified for *E. coli* IspE ^23^ where also a similar domain arrangement to the other *E. coli* IspE crystal structure is evident ^14,18^. Specifically, the newly found crystals have the space group C2 and diffract at 1.5 Å resolution (refinement statistics can be found in Table S2). In this monoclinic crystal form, the active site pocket is fully accessible for ligand binding as the binding sites of the two subunits were positioned in opposite directions (180° rotation from each other). This configuration created a fully accessible pocket for ligand binding allowing the use of soaking or co-crystallization (Fig. 3). We obtained complex structure of *E*.*coli* IspE with CDP-ME and the unhydrolyzable ATP analog, Adenylyl-imidodiphosphate (ANP), for which a clear density was observed for only two phosphates and not the terminal phosphate, most probably because of the flexibility of the triphosphate. Consequently, we placed ADP in the pocket instead of ANP. The binding of both molecules closely resembles what has been described previously for *E. coli* IspE ^14^.

**Figure 3:**
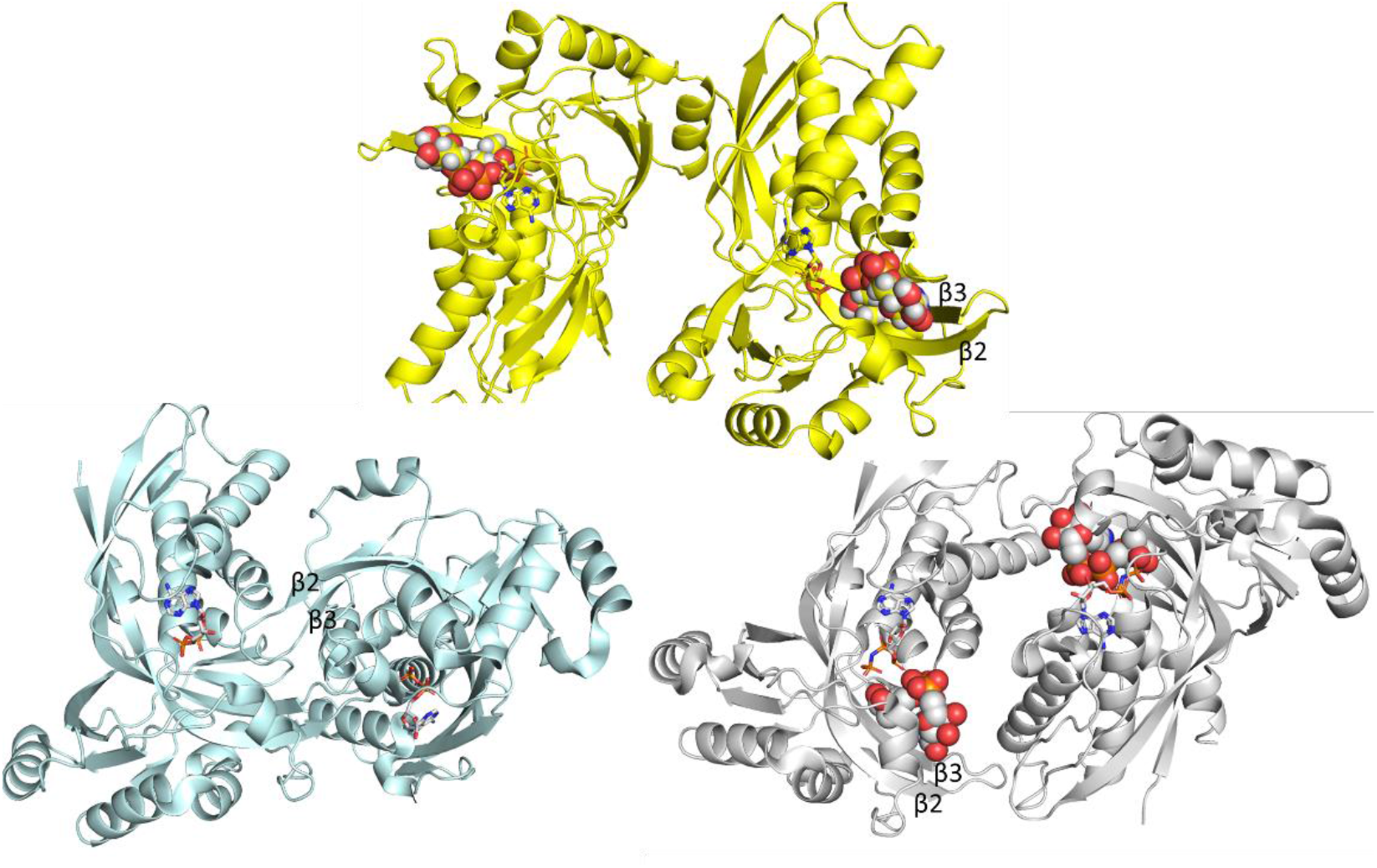
Comparison of *Escherichia coli* IspE crystal conformations in the ASU. Differences between the two monoclinic crystal forms and the triclininc are highlighted. PDB: 8QC7 (monoclinic C121), obtained in this study, is shown in yellow, PDB: 1OJ4 (monoclinic, P 1 21 1) is shown in gray, and PDB: 2WW4 (triclinic P1) is shown in light blue. Molecule A from all crystals is presented in the same orientation to illustrate substrate (CDP-ME, shown as spheres) availability/blockage. ANP is shown as sticks.

The total solvent-accessible area in the new crystal form is 25.127 Å^2^ and the buried interface area is 97.0 Å (3%), compared to 1167.9 Å (23%) in the triclinic structure and 568.7 Å^2^ (12%) in the previously reported monoclinic space group *P*21 (calculated using PISA server). Figure 3 shows the different crystal forms identified for *E*.*coli* IspE in comparison to those obtained in this study.

### 2.3 Biochemical evaluation of IspE substrate-competitive inhibitors

In the search for inhibitors against IspE, it is possible either to target the ATP or the substrate-binding sites. Kinase inhibitors are mostly designed to target the ATP–binding site ^25,26^. In IspE kinase however, this pocket is rather shallow and mostly solvent-exposed. Over the years, our group attempted to address this pocket. We, recently, performed a virtual screening on the ATP binding pocket followed by an SAR study ^27^, and in the past Tang et al. used a library of existing GHMP kinases inhibitors ^28^. Unfortunately, both attempts were not very successful in developing selective IspE inhibitors targeting the kinase ATP pocket.

On the other hand, the substrate-binding pocket and subpockets have been successfully targeted ^22,29,30^, and the work published in 2007 led to the identification of compound (±)-**1** ^29,31^, Figure 4. Brenk group also extensively focused on the study of IspE inhibitors and they reported an *in silico* and high-throughput (HTS) approaches that identified inhibitors in the high micromolar range ^32^. It is evident that major efforts are necessary to optimize the existing inhibitors as well as to identify novel hit molecules and, to do so further insights about the binding mode could be very beneficial. Here, we used compound (±)-**1** as a reference inhibitor due to its high potency and selectivity against IspE. As shown in Fig. 4, this structure consists of a central cytosine scaffold bearing a tetrahydrothiophene ring, as ribose analogue, bridged via a triple bond to a cyclopropyl sulfonamide. Aiming to identify determinants for IspE inhibition by (±)-**1**, we conducted a focused structure-activity relationship (SAR) study that separately targeted each component of the molecule.

**Figure 4:**
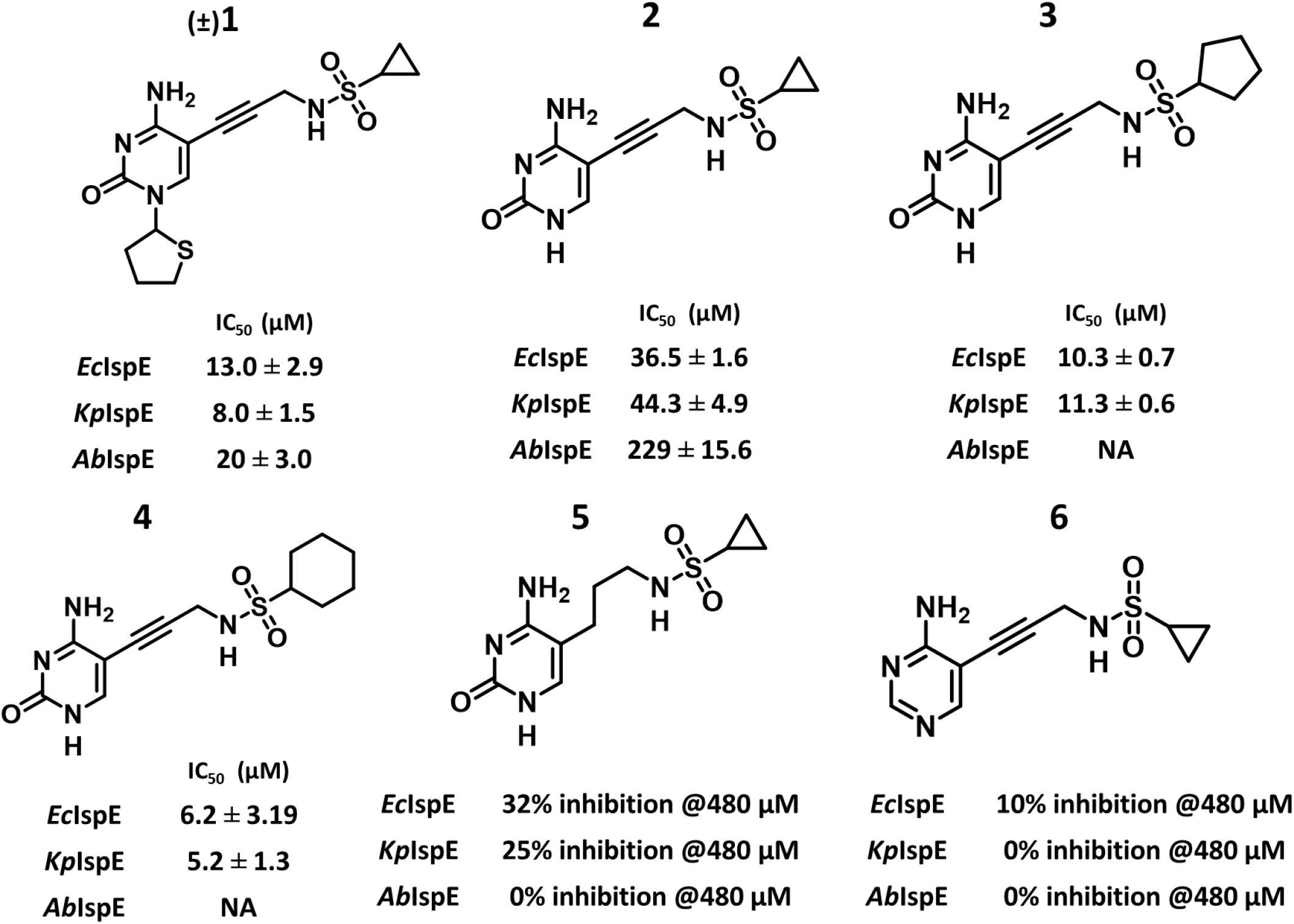
Biochemical evaluation of compounds (1–6). The half-maximal inhibitory concentration (IC50) on *Escherichia coli* IspE (*Ec*IspE), *Klebsiella pneumoniae* IspE (*Kp*IspE) and *Acinetobacter baumannii* IspE (*Ab*IspE) are depicted. The values presented are average of duplicate measurements. NA: not available; no dose-response curve was obtained to calculate the IC50 value.

The designed and synthesized subset of compounds (**1**–**6**) were simultaneously tested *in vitro* on the three IspE homologs studied in this work (Figure 4). Interestingly, we found that all the compounds showed a similar inhibition pattern against *E. coli* IspE and *K. pneumoniae* IspE, with **5** and **6** being inactive. As expected, the biggest difference is found against *A. baumannii IspE* where, only compound **(±)-**1 is still active.The three homologs exhibited an inhibition pattern that correlates with their structural similarities and differences in the binding site, indicating the need for different strategies in designing inhibitors targeting each of these enzyme homologs.

Next, we selected the most potent and soluble compounds for co-crystallization to further confirm their binding mode and guide our optimization strategies. Although we could obtain high-resolution complex structures with *E. coli* IspE, the molecular packing of *K. pneumoniae* IspE prevents substrate -and inhibitors designed to bind to the substrate pocket-binding. Nevertheless, due to their high similarity, we propose that the binding mode can be translated from *E. coli* to *K. pneumoniae*. Encouragingly, this assumption is also experimentally supported by the comparable IC_50_ values found. On the other hand, all our efforts have failed in obtaining a complex structure with *A. baumannii* IspE, due to its high solubility, which means that high concentrations are required to drive crystallization. In addition, because *A. baumannii* IspE has 13% lysine residues in its sequence, mostly on the surface, we tried lysine methylation which resulted in poor-quality crystals.

### 2.4 *E. coli* IspE-inhibitor complex structures

Although the new monoclinic *E. coli* IspE crystals allow soaking experiments, high concentrations of the inhibitor is required in order to exchange the substrate by the inhibitor; therefore, the inhibitors have to be sufficiently soluble. Fortunately, the substrates of the MEP pathway, especially those of the IspE enzyme, are highly polar due to their charged di- and triphosphate groups^33^. As mentioned earlier, for this study our choice was directed towards **(±)**-**1** and its derivatives. In all the complex structures obtained with *E. coli* IspE and compounds **1** (PDB: 8QCC), **2** (PDB: 8QCN), and **3** (PDB: 8QCO), a consistent interaction pattern was observed. The central cytosine in the inhibitors is positioned between Tyr25 and Phe185, leading to the formation of hydrogen bonds with both the backbone and side chain of His26, similar to the interaction observed with the cytosine ring in CDP-ME, (Fig. 5). In the structure with compound **(±)-1**, the tetrahydrothiophene ring overlaps with the ribose moiety of CDP-ME, where it occupies a central position between Tyr25 and Pro182, forming a pseudo-π sandwich (Fig. 5A). While these residues are conserved in the *K. pneumoniae* IspE structure, Pro182 is replaced by a Gln174 in *A. baumannii* IspE, which means a different type of interaction will have to take place in this position. Additionally, the sulfur atom of the tetrahydrothiophene ring could potentially point to form hydrogen bonds with either Thr181 or Tyr25. The alkyne linker guides the sulfonamide moiety into the notably polar phosphate binding region situated between the substrate and the ATP binding sites. The nitrogen atom of the sulfonamide forms a hydrogen bond with the side chain of Asp141. Furthermore, the sulfone group, in relation to the NH group, adopts a staggered conformation, facilitating hydrogen bond interactions with the side chain of Asn12 and Lys10. In compounds **1, 2** and **3**, the orientation of the cyclopropyl and cyclopentyl rings is directed towards a small hydrophobic pocket delineated by Leu15, Leu28, and Phe32. This hydrophobic region partially contributes to binding the methyl group of the substrate, CDP-ME. Our experimental findings confirm that these compounds are indeed binding to the substrate-binding pocket as intended, forming interactions similar to those observed with the native substrate.

**Figure 5:**
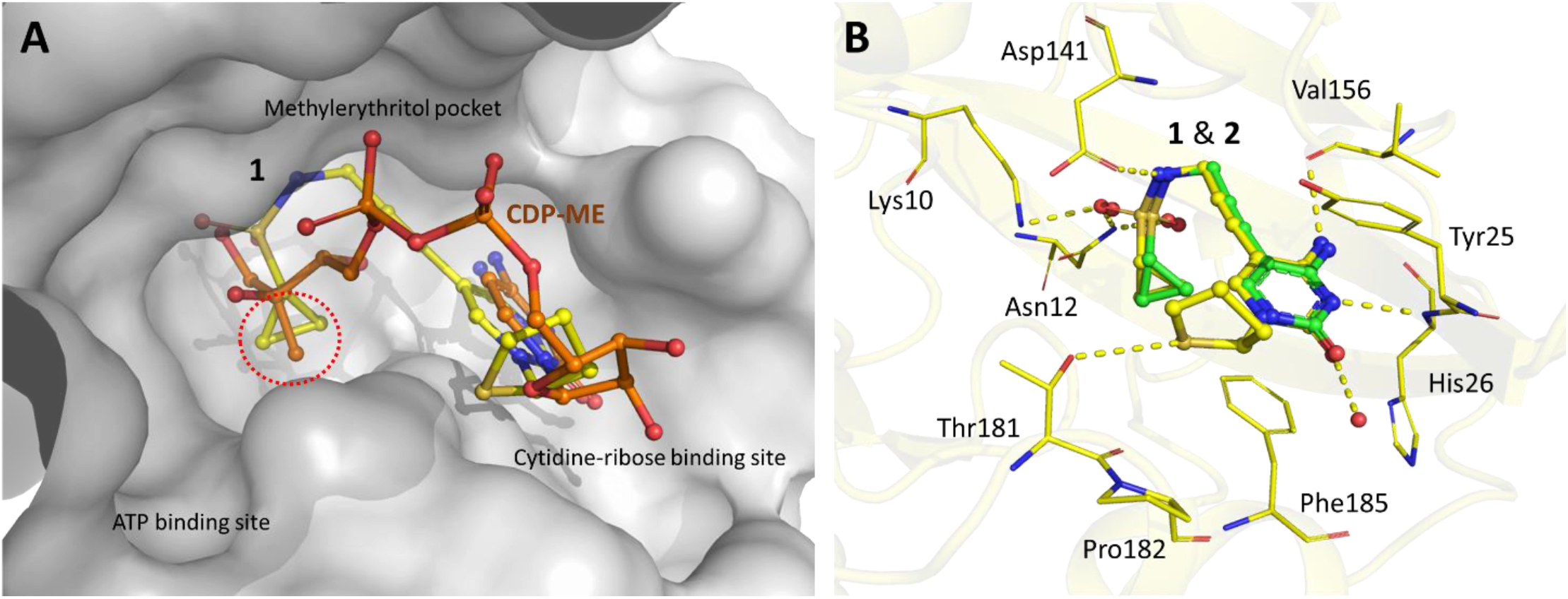
Binding of compound (±)-1 and 2 in the substrate binding site of *Escherichia coli* IspE: **A:** Surface representation of*E. coli* IspE active site, showing an overlay between the CDP-ME and the competitive inhibitor **(±)-1** in the substrate pocket. The active site of IspE is composed of three main pockets: the cytidine binding pocket with the ribose sub-pocket, the methylerythritol (ME) binding site featuring a small hydrophobic cavity (highlighted by a red circle), and the ATP binding pocket. **B:** Superimposition of experimental structures of *E. coli* IspE co-crystallized with compounds **(±)1** in yellow and **2** in green showing binding interactions discussed in the text. Molecular surface representations performed using PyMol. Color code: C: green and yellow; O: red; N: blue; S: yellow.

Compound **2** was designed to experimentally evaluate the importance of the tetrahydrothiophene ring. Interestingly, it demonstrated an almost three-fold decrease in activity with IC_50_ values against *E. coli* IspE and *K. pneumoniae* IspE respectively, and a 10-fold reduction against *A. baumannii* IspE . The complex structure shows that this drop in activity was due to the loss of the tetrahydrothiophene ring`s hydrophobic interaction (stacked between Tyr25 and Pro182) and of a potential hydrogen bond of the S atom with Thr181 or Tyr25 (Fig. 5B). Despite the dramatic decrease in activity especially against *A. baumannii* IspE, the compound still retained some activity against the other homologs, indicating that the rest of the molecule still significantly contributes to binding.

Aiming to test the optimum binding occupancy of the small hydrophobic pocket, we designed and synthesized compounds **3** and **4** bearing a five and six member ring, respectively in place of the cyclopropyl of **(±)-1**. The analysis of the co-crystalization with *E. coli* and *K. pneumoniae* IspE showed that both compounds have regained the lost activity in compound **2** and slightly improved the affinity compared to **(±)-1**. Compound **4** with the six-membered cyclohexyl ring further improved the inhibitory activity compared to compound **3** with the five-membered cyclopentyl ring. Theoretical and experimental studies have previously shown that occupation of hydrophobic pockets in a protein target is the main contributor to the binding affinity of the ligand ^34^. In IspE, it was found that this hydrophobic pocket, originally accommodating the methyl group of the methylerythritol moiety, is notably more spacious than initially anticipated. Therefore, the appropriate filling of this sub-pocket would result in a significant gain in binding free enthalpy ^35^. The volume of this pocket measures roughly around 180 Å^3^ in *E. coli IspE* and *K. pneumoniae* IspE, while it is only 102 Å^3^ in *A. baumannii* IspE. Prior studies indicated a significant relationship between its occupancy and activity of the ligands targeting the substrate pocket ^30^.

Although we could not obtain a complex structure with compound **4**, due to solubility issues, a co-crystal structure of compound **3** with *E. coli* IspE was obtained, which showed that the cyclopentyl ring indeed occupies the deep hydrophobic pocket surrounded by nonpolar residues, Ile15, Ile17, Leu28 and Ph185 (Fig. 6). This hydrophobic interaction exhibited sufficient strength for these two compounds to possess higher affinities than compound **(±)-1**. This factor should be taken into account when considering an extension from this subpocket to the adjacent phosphate-binding pocket.

**Figure 6:**
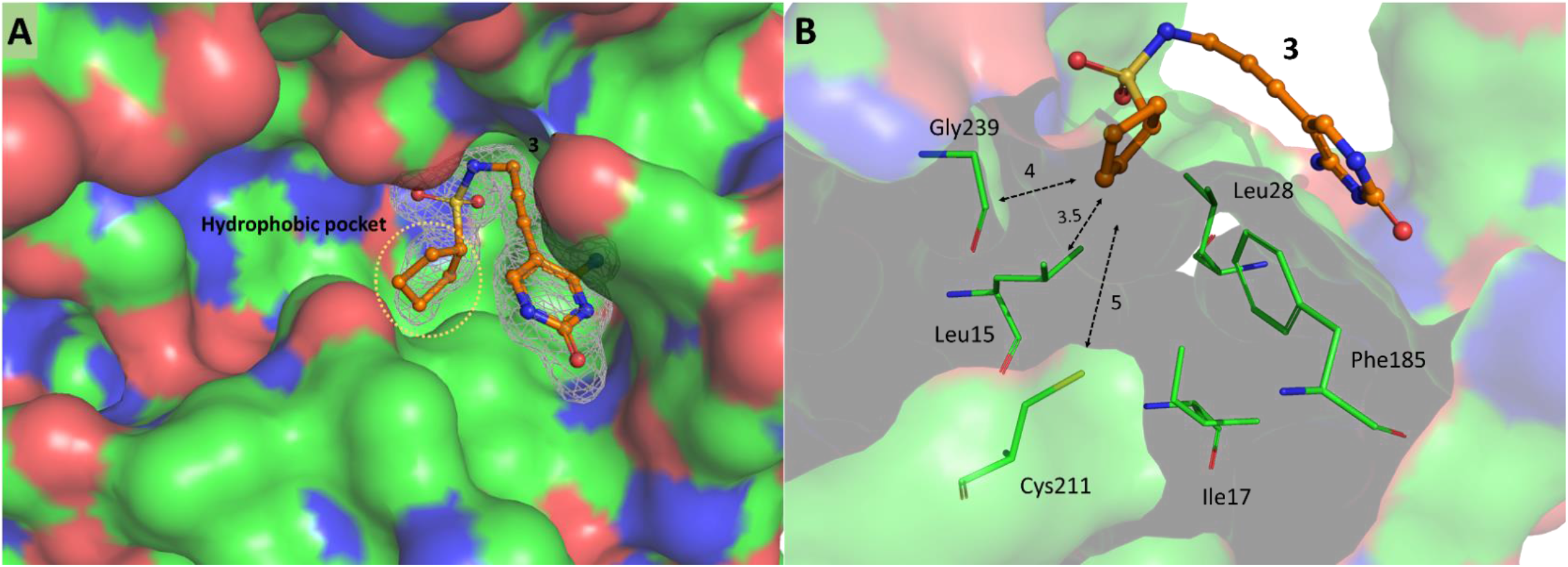
Binding of ligand 3 to *Escherichia coli* IspE (PDB: 8QCO): **A:** Molecular surface representation of *E. coli* IspE showing the hydrophobic pocket occupied by the five-membered hydrocarbon ring from the ligand. **B:** Close-up of the small, hydrophobic pocket showing residues of the receptor surrounding the ligand, distance to the closest residues in the hydrophobic pocket are shown in Ångstrom. Color code: C: green; O: red; N: blue; S: yellow.

### 2.5 *A. baumannii* IspE needs to be targeted using a tailored strategy

Among the subset of compounds synthesized and tested, only **(±)-1** showed inhibition against IspE from *A. baumannii* comparable to the other tested homologs On the contrary, the inhibition values for **2, 3** and **4**, are substantially different. This low activity might also be the reason for the absence of X-ray crystal structure from this homolog To overcome this bottleneck, we used sequence alignment and homology modeling to rationalize this and to structurally showcase the differences.

The Swiss-Model of *A. baumannii* IspE (Uniprot accession code: B2HUM5) was aligned with the *E. coli* IspE structure in MOE with an overall RMSD of 1.118 Å ^36, 37^. Fig. 7, shows the structural differences between the two homologs within the CDP-ME binding site.

**Figure 7:**
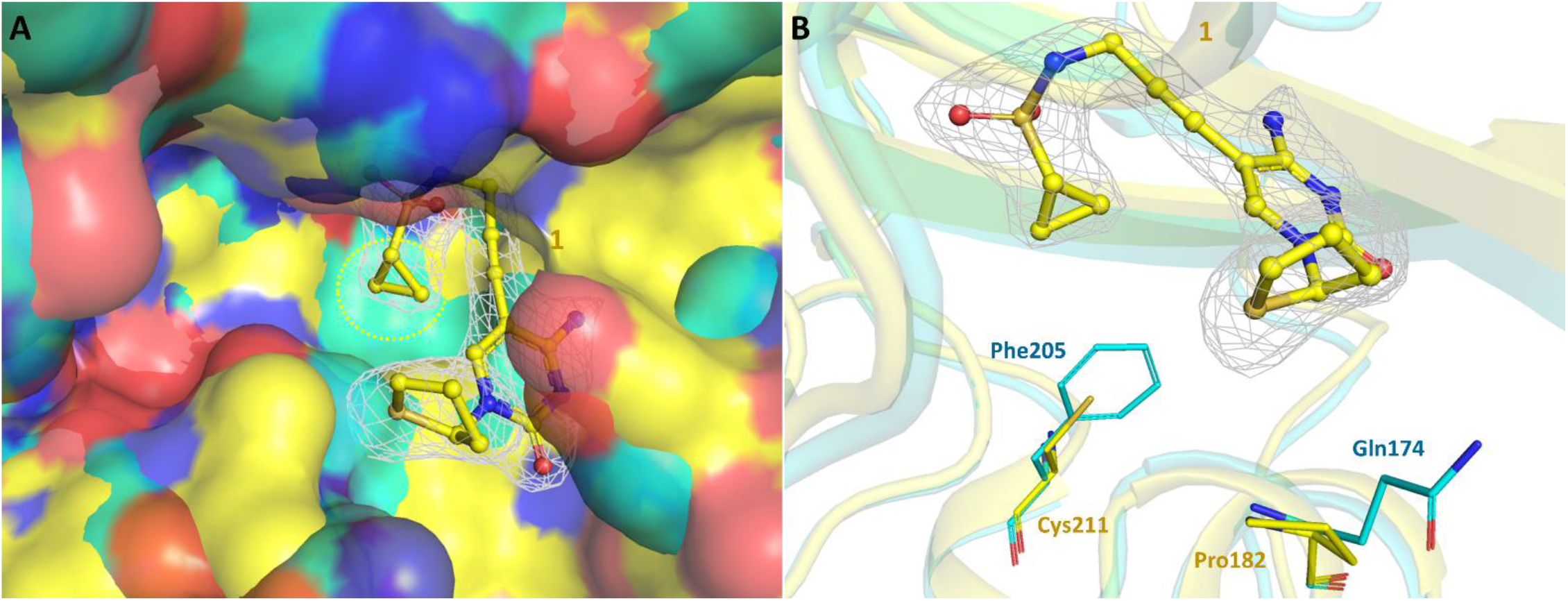
*Escherichia coli* IspE binding to (±)1 superimposed with homology modeled structure of *Acinetobacter baumannii* IspE. *E. coli* IspE in yellow superimposed with *A. baumannii* IspE in cyan. **A:** molecular surface representation of the *A. baumannii* IspE showing the tighter hydrophobic pocket targeted by the cyclopropane in the ligand. **B:** Showing a close-up of the main difference in the active site between the two homologs. In which a cysteine (Cys211) in *E. coli IspE* is replaced by a phenylalanine (Phe205) in *A. baumannii* IspE. Color code: C: cyan and yellow; O: in red; N: in blue; S: yellow.

Compounds **2**-**4** were docked into the CDP-ME binding site of *E. coli* IspE (experimental crystal structure) and *A. baumannii* IspE (Swiss model) resulting in highly similar binding modes. The docking scores are in line with experimental inhibition data showing a slightly lower predicted binding energy of **2** compared with **3** and **4** (Table S2). We attribute this to the reduced ability of the cyclopropyl ring to interact with the small hydrophobic pocket at the back of the binding site.

On the other hand, when looking at Fig. 8A, B, and C, we observe significant differences of the binding mode of these compounds in the *A. baumannii* IspE structure. In particular, for **2** (Fig. 8A), there is loss of the interaction of the cytosine carbonyl with His25 as well as the loss of the interaction between the sulfonamide oxygen and the side chain of Asn11 (when compared to the *E. coli* IspE binding mode). Similarly, for **3** (Fig. 8B), we observe the loss of hydrogen bonding interactions between the cytosine ring and the backbone of Ile48 with the formation of an internal hydrogen bond instead. Finally, for **4** (Fig. 8C), the interactions of the cytosine ring with the backbone are maintained, however, in the docked pose, the sulfonamide no longer interacts with the side chains of Asn11 and Asp133 leading to a loss in predicted binding energy relative to the *E. coli* IspE structure. Taking the inhibition values in mind, there seem to be no side chain movements enough to restore those lost interactions in the *A. baumannii* homolog.

**Figure 8:**
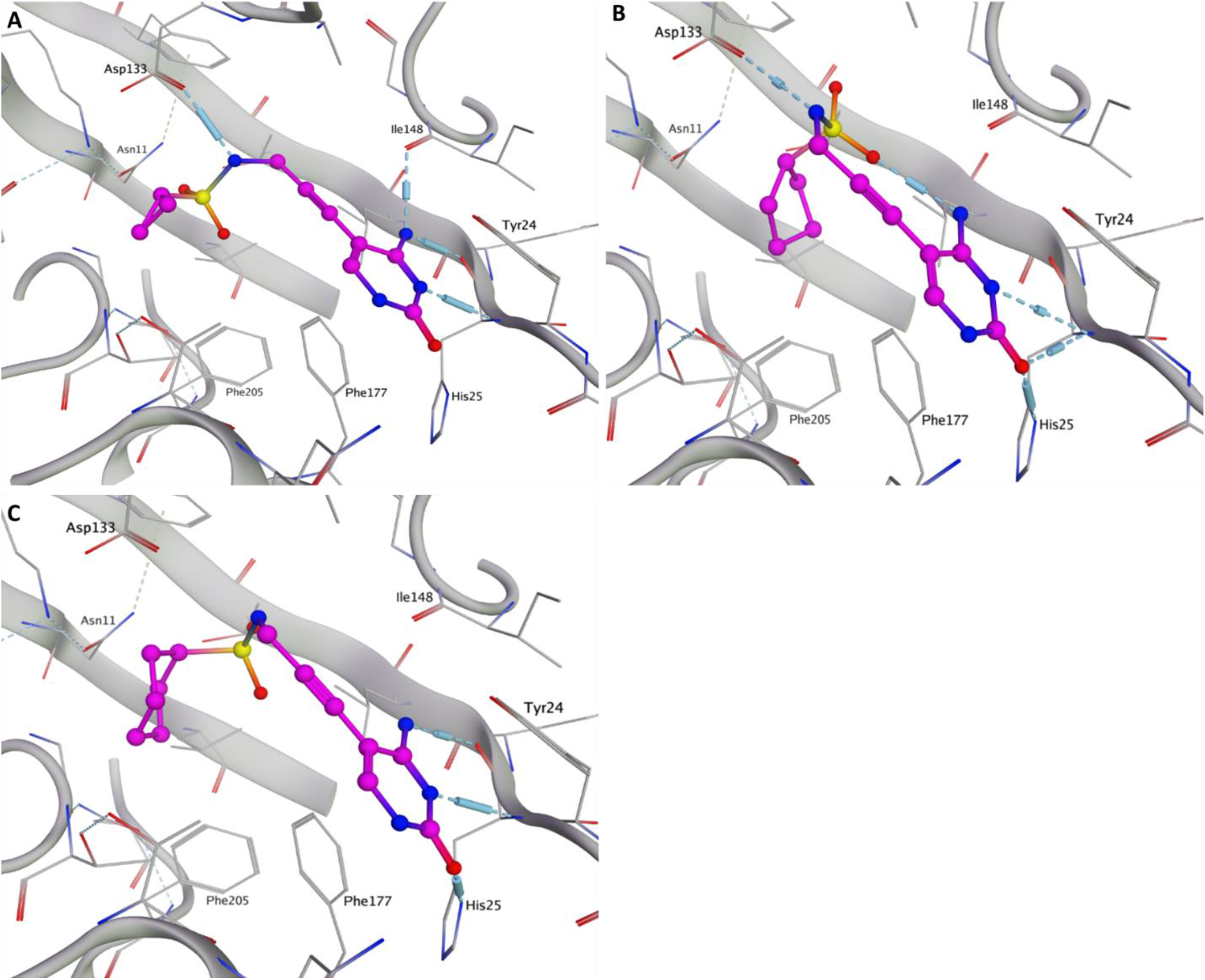
Docking of compounds 2,3 and 4 in *A. baumannii* IspE. **A**: Best docked pose of **2** (in magenta) in the *A. baumannii* IspE SwissModel structure (in gray) showing loss of the interaction of the cytosine carbonyl with His25 and the sulfonamide oxygen with the side chain of Asn11. **B:** Best docked pose of **3** (in magenta) in the *A. baumannii* IspE SwissModel structure (in gray) showing loss of hydrogen bonding interactions between the cytosine ring and the backbone of Ile48 and the formation of an internal hydrogen bond instead. **C:** Best docked pose of **4** (in magenta) in the *A. baumannii* IspE SwissModel structure (in gray) showing the shift of the molecule binding, the sulfonamide losses interacttion with the side chains of Asn11 and Asp133. Relevant residues are labelled in each picture. Protein residues and ligands shown as sticks, C: gray and magenta; O: red; N: blue; S: yellow.

Replacing of the Cys211 in *E. coli* IspE with Phe205 in *A. baumannii* IspE appear to reduce the size of a small hydrophobic pocket. In *E. coli* IspE, this pocket accommodates the cyclopentyl ring of compound **3**. However, the smaller size *in A. baumannii* IspE causes a shift of the compound and a concomitant loss of the important hydrogen bonds with either the cytosine ring or the sulfonamide moiety (Fig. 8A). This explains the observed decrease in experimental activity of these compounds against *A. baumannii* IspE. The presence of glutamine instead of proline in *A. baumannii* IspE creates an opportunity on the opposite side of the pocket. We can design the compound to form a hydrogen bond with this glutamine residue, potentially improving binding affinity. Furthermore, the 3-membered ring seems to be optimal for interaction within this pocket. Keeping this core structure while expanding the molecule towards the adjacent ATP binding site could be a promising strategy. This expansion might lead to additional interactions with the protein, potentially improving binding and activity. By combining these approaches, we can potentially develop more potent compounds against *A. baumannii* IspE.

On the other hand, for the homologs from *E.coli* and *K. pneumoniae*, the hydrophobic pocket can be further utilized by incorporating a functional group that forms a covalent bond with the cysteine within the pocket. This stronger interaction could enhance the compound’s effectiveness.

Few additional findings are worth mentioning; we tested the importance of the alkyne linker by synthesizing compound **5** with a flexible linker. This resulted in a complete loss of activity across all homologs suggesting the important role of this alkyne linker to direct the cycloalkyl rings into the hydrophobic pocket. The study also highlighted the challenge of cytosine replacement without compromising ligand affinity, a contrast to the numerous adenine substitutions seen in kinase inhibitors. The replacement of cytosine with just an amino pyrimidine ring resulted in a significant reduction in activity as observed in compound **6**. It is also worth noting here that the most active compound (**4**) has an IC_50_ value of 5.2 µM, which leaves room for further optimization. Although, the exclusion of the thiophene ring is a major step in the simplifying these derivatives, shortening the synthesis and allowing for higher final yields. These preliminary findings will guide our future synthesis in the development of novel IspE inhibitors.

## 3 CONCLUSIONS

Our study aimed to characterize IspE kinase homologs, explore the substrate pocket and study inhibitor binding modes. To achieve this, we recombinantly overexpressed and purified the proteins from *E. coli* IspE, *K. pneumoniae* IspE, and *A. baumannii* IspE, established biochemical assays, and obtained complex structures with inhibitors. While *E. coli* and *K. pneumoniae* IspE exhibited similar kinetics, consistent with previous reports. In addition to showing lower sequence identity, *A. baumannii* IspE kinetic values highlighted its divergence. These values were used in optimizing the assay conditions for each homolog, setting the stage for high-throughput screening in search of inhibitors.

We presented the first crystal structures of *K. pneumoniae* IspE in complex with the non-hydrolysable ATP analog. The 3D-structural characteristics and kinetic parameters of *K. pneumoniae* IspE closely resemble those of *E. coli* IspE. Therefore, we suggest that they can be employed interchangeably, particularly when encountering challenges in obtaining and/or crystallizing either enzyme. We obtained a new crystalline form for *E. coli* IspE, which allows ligand binding during soaking. We successfully used this condition to obtain three new complex structures of *E. coli* IspE with inhibitors, which allowed us to gain deeper understanding of the role a the methylerythritol hydrophobic subpocket in determining activity of compounds targeting the substrate binding site. This pocket is relatively bigger in *E. coli* IspE and *K. pneumoniae* IspE and improved compounds activity was observed by its proper filling. Covalent targeting of a cysteine residue deep in the pocket might also be considered for further improvements of this compounds class. For *A. baumannii* IspE, the attention could be drawn to the other side of the pocket where a glutamine is replacing a proline, that is important for interaction with CDP-ME and ligands designed to target the substrate pocket.

Our findings emphasize the nuanced nature of IspE kinase inhibition, guided by subtle structural variations among homologs and offer valuable insights into the development of potent and selective inhibitors for IspE kinase homologs, contributing to the broader goal of combatting pathogenic microorganisms reliant on the MEP pathway.

## 4 MATERIALS AND METHODS

### Protein preparation

We transformed pET28a plasmids containing our constructs into electrocompetent BL21 (DE3) *E. coli* cells. The transformed cells were subsequently cultured on LB-Agar supplemented with 25 µg/mL kanamycin and allowed to incubate overnight at a temperature of 37° C. We then transferred the selected colonies into LB medium supplemented with 25 µg/mL kanamycin, at 37 °C until OD_600_ 0.3–0.6 for *E. coli* IspE and *K. pneumoniae* IspE, and 0.8 for *A. baumannii* IspE. We induced overexpression with 1 mM IPTG/18° C/180 RPM/overnight. The harvested cells were then disrupted before subsequently purifying the proteins. We first performed affinity chromatography using HisTrap HP 5 mL, then removed tags using TEV protease (1:20). The output from the reverse IMAC column was injected into a 7 mL loop of the ÄKTA pure system on an S200 SEC column. Peak fractions were combined and concentrated in the respective storage buffer for each protein. Finally, we performed SDS-PAGE gel electrophoresis to confirm the presence and purity of the purified IspE homologs at each purification step (Fig. S2).

### Gel filtration analysis

Purified samples of *E. coli* IspE, *K. pneumoniae* IspE and *A. baumannii* IspE were prepared for the gel filtration analysis. To determine the molecular weights of our proteins, standard proteins of known molecular weights were used to calibrate the gel filtration column. Calibration standards included proteins ranging from 670 kDa to 1.35 kDa. A defined volume of the prepared *K. pneumoniae* IspE sample was injected into the calibrated gel filtration column. The protein was detected using UV spectroscopy at wavelength of 280 nm. The elution profile of IspE was recorded based on the absorbance signals from the UV detector. The molecular weight of IspE was estimated by comparing its elution volume to the elution volumes of the calibration standards. A calibration curve was constructed using the known molecular weights of the standards (Fig. S2).

### Enzymatic assay

To assess IspE activity and its kinetic parameters, we adapted a coupled spectrophotometric enzyme assay based on a previously described protocol ^19^. The analysis was carried out at room temperature (RT) over a 30-minute period. Enzymes and substrate concentrations were adapted for each homolog to ensure a suitable reaction velocity and linear kinetics. *E. coli IspE* in concentration of 200 nM, *K. pneumoniae* IspE of150 nM and *A. baumannii* IspE of 100 nM. For the kinetic assays, varying concentrations of the cofactor ATP and the substrate CDP-ME were employed. When one of these was held constant, ATP was maintained at 1 mM, and CDP-ME at 2 mM. For the determination of the IC_50_ values of specific inhibitors, the percent of inhibition was calculated from a linear curve of initial velocities of the enzymes with different concentrations of the compounds. Kinetic parameters and IC_50_ values of inhibitors were calculated using nonlinear regression analysis (OriginPro2023) (Fig. S3).

### Crystallization

To determine the optimal crystallization condition of IspE homologs in the study, screening trials were carried out using commercially available crystallization kits from NeXtal. Plates were incubated at 18 °C and checked for crystals every three days. Repetitive optimizations were performed for the best identified crystal conditions. ***K. pneumoniae* IspE**: the structure was obtained after incubating 15 mg/mL *K. pneumoniae IspE* with 5 mM of each of the AMP-PNP, CDP-ME and MgCl_2_. Crystals were set in vapor diffusion chambers with a reservoir solution consisting of 1000 mM Sodium citrate, 100 mM HEPES pH 7.0. ***E. coli* IspE**:7.5mg/mL protein was used to grow crystals in vapor diffusion chambers with a reservoir solution consisting of 25% PEG 4k; 200mM MgCl_2_; 100mM MES pH 6.5. The crystals were then soaked for 2 days with 5–10 mM compound. The best crystals were then mounted with 32% of glycerol (as a cryo-protectant) and data collection were carried out using beamtimes provided by the SLS and ESRF.

### X-ray data collection and structures determination

The structure of the new crystal form of *E. coli* IspE and *K. pneumoniae* IspE was determined through molecular replacement using Phaser.MR ^38^ within the Phenix software suite ^39^. The primary model used for the search was *E. coli* IspE (PDB ID: 1OJ4) ^14^. Initially, data reduction and alignment were carried out using AIMLESS ^40^ within CCP4i ^41^. Following this, manual interpretation of the electron density maps was conducted using COOT ^42^. The 3D structures were refined iteratively using Phenix refine (43). Structures were validated in the PDB validation server before deposition. For visual representation, all figures were generated using PyMoL ^43^. Comprehensive statistics regarding data collection and the refinement process can be located in Table S2.

### Compounds synthesis

We sourced chemicals from commercial suppliers without further purification for our chemical synthesis. Chemical yield measurements are based on purified compounds, and were not optimized. Reaction progress was monitored with TLC silica gel sheets and UV visualization. Column chromatography used an automated CombiFlash® Rf system with RediSepRf silica columns. Preparative RP-HPLC was performed with a Thermo Fisher Scientific UltiMate 3000 system and a nucleodur® C18 Gravity column. ^1^H and ^13^C NMR spectra were acquired using Bruker Avance Neo 500 MHz or Bruker Fourier 300 instruments. Chemical shifts were referenced against solvent peaks, and splitting patterns were noted. Low-resolution mass analysis and purity checks used an Ultimate 3000-MSQ LCMS system. Compounds for biological assays had ≥ 95% purity. High-resolution mass spectra were obtained using a ThermoFisher Scientific Q Exactive Focus system.. For more detailed chemical and analytical methods, please refer to the supplementary materials.

### Computational methods

Docking studies for IspE inhibitors to *E. coli* IspE and *A. baumannii* IspE were conducted using MOE v2020.09. The ligand preparation involved creating 2D structures of compounds and subjecting them to energy minimization. Protein structures were prepared for *E. coli* IspE and *A. baumannii* IspE through force-field application, hydrogen assignment, and energy minimization. Binding sites were selected based on proximity to ligands. Molecular docking was performed using MOE’s Dock module, utilizing the co-crystalized ligand as a template and specific receptor structures. Scoring functions were applied to assess docking results. Detailed computational methods can be found in the supplementary.

## Supporting information

Hamid, et.al, IspE as a Drug Target- Supplementary

## Acknowledgments

We acknowledge the European Synchrotron Radiation Facility (ESRF), Grenoble, France for provision of beamline ID23-1 and ID30B. We acknowledge the Paul Scherrer Institute, Villigen, Switzerland for the provision of synchrotron radiation beamtime at beamline X06DA-PXIII and X06SA-PXI of the SLS.

The authors would also like to thank Boris Illarionov for providing the substrate CDP-ME and Paula Kramer for helping with the computational work.

## Author contributions

A.K.H.H., R.H., conceived the study. R.H. wrote the main manuscript text and prepared all figures. D.W. and E.D. synthesized the compounds for bioassays. D.A. performed the bioassay, A.L. performed the docking study. M.M.H designed and supervised the medicinal chemistry work. All authors proofread the manuscript. A.K.H.H. supervised the project.

## Funding

The Helmholtz Association’s Initiative and Networking Fund and the Schlumberger Foundation faculty for the future (FFTF) funded this work.

## Competing interests

The authors declare no competing interests.

## Data availability

The authors confirm that the data supporting the findings of this study are available within the article [and/or] its supplementary materials.

## Declaration of generative AI and AI-assisted technologies in the writing process

During the preparation of this work the author(s) used ChatGPT-3.5 in order to improve language and readability. After using this tool, the author(s) reviewed and edited the content as needed and take(s) full responsibility for the content of the publication.

## Notes

### Competing Interest Statement

The authors have declared no competing interest.

