## Supplementary material for "IspE Kinase as an Anti-infective Target: Role of a Hydrophobic Pocket in Inhibitor Binding": Hamid, et.al, IspE as a Drug Target- Supplementary

### IspE as a Drug Target: Reading the Role of a Hydrophobic Pocket in Inhibitor Binding


| | |
|---|---|
| Figure S1: Michaelis-Menten kinetics analysis, $K_m$ values of *E. coli* IspE, *K. pneumoniae* IspE and *A. baumannii* IspE | 2 |
| Figure S2: Sequence alignment of the IspE from *K. pneumoniae*, *E. coli and A. baumannii* | 3 |
| Figure S3: Gel filtration calibration for molecular weights calculation and SDS-PAGE 12% electrophoresis | 4 |
| Figure S3: Inhibitory dose-response curves | 5 |
| Table S1: Data collection and refinement statistics | 6 |
| Scheme S1: Compounds synthesis | 7 |
| Table S2: Docking scores of the three compounds docked to *E. coli* IspE and *A. baumannii* IspE | 22 |

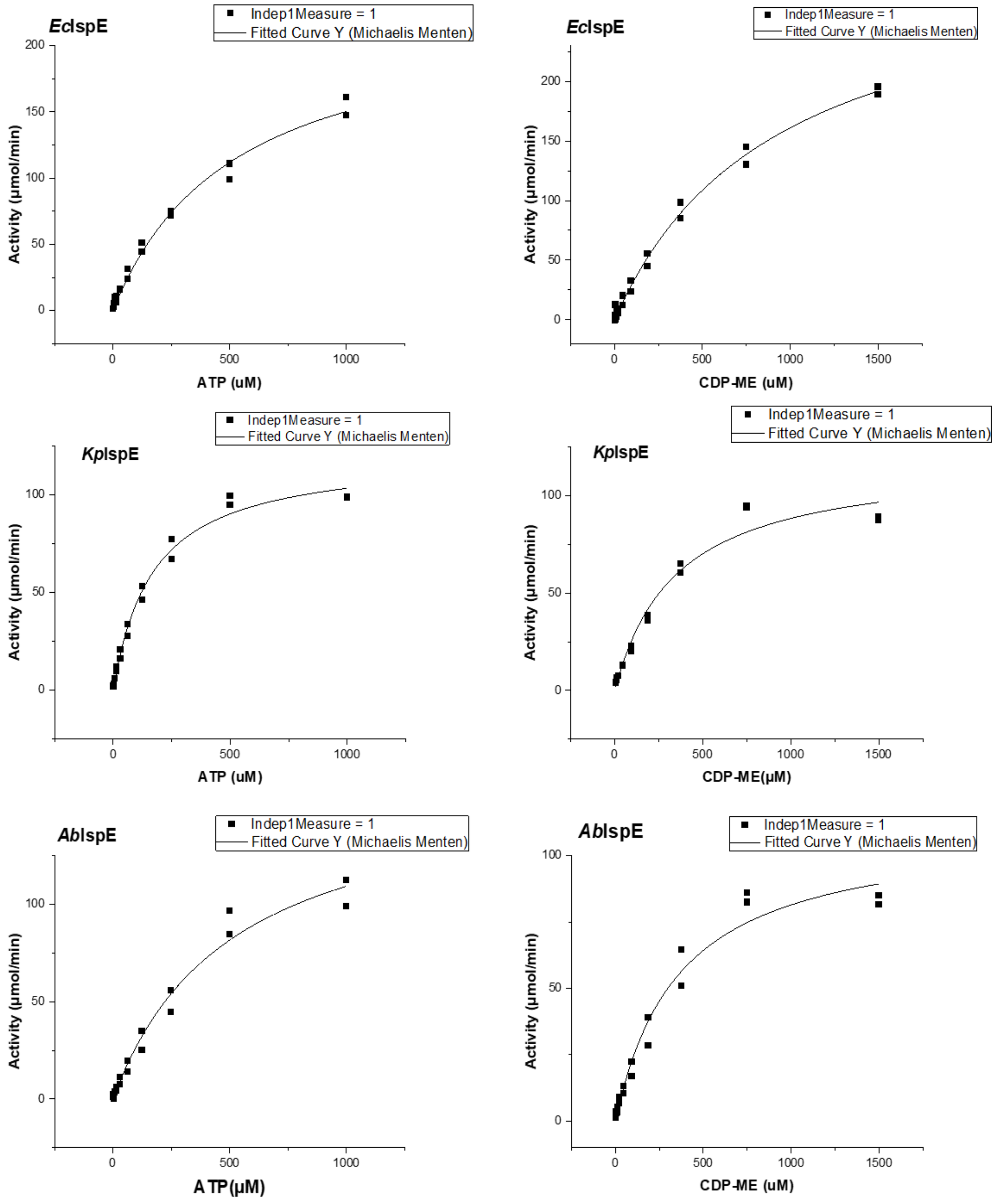


**Figure S1: Michaelis-Menten kinetics analysis, $K_m$ values of *E. coli* IspE, *K. pneumoniae* IspE and *A. baumannii* IspE enzymes**, analysis was conducted by varying concentrations of ATP and CDP-ME, enzyme concentration used: *E. coli* IspE (200nM), *K. pneumoniae* IspE (150nM) and *A. baumannii* IspE (100nM). When a substrate or cofactor was kept constant, a concentration of 1mM was used for ATP and 1.5 mM for CDP-ME.

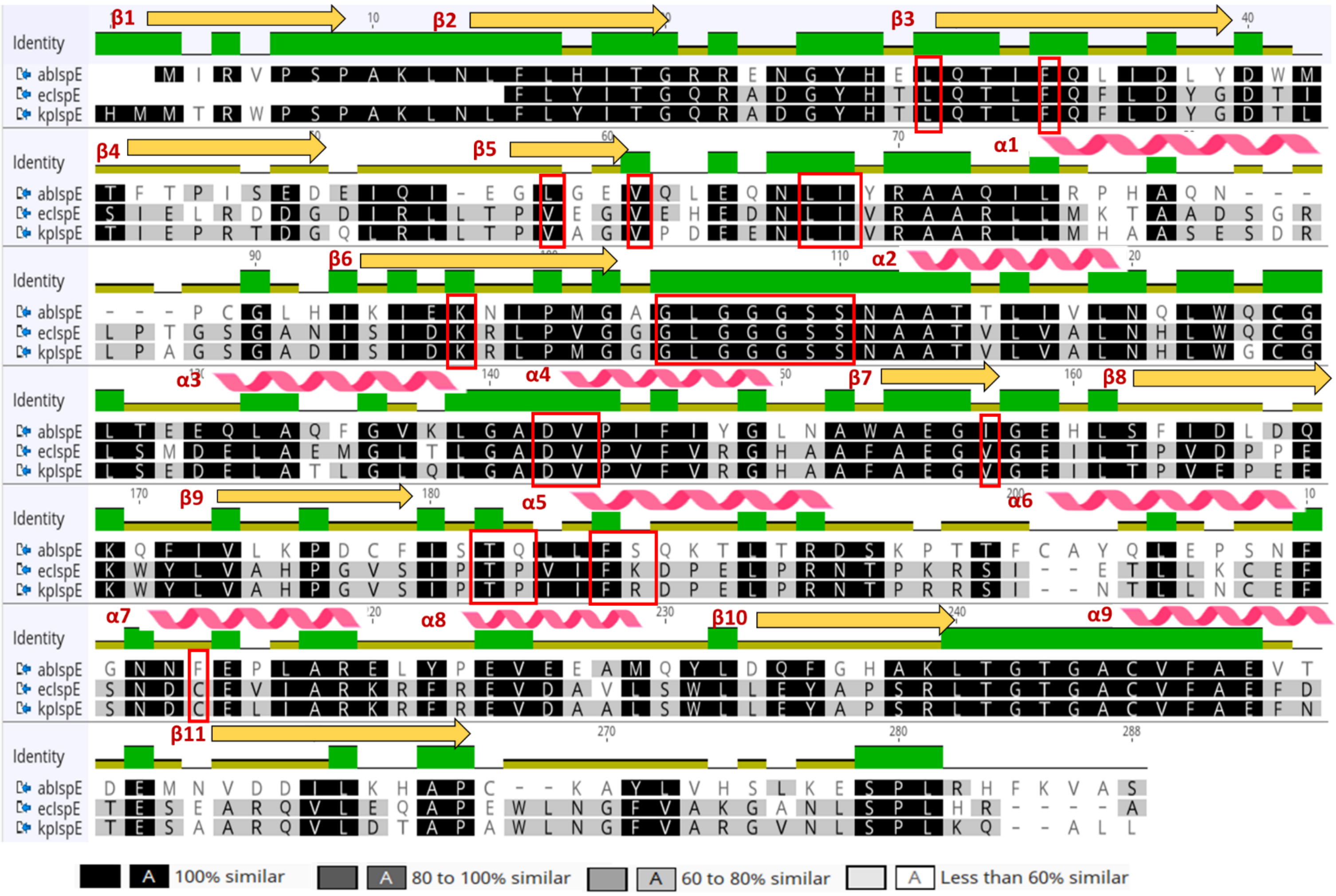


| | abIspE | ecIspE | kpIspE |
|---|---|---|---|
| abIspE | | 39.051% | 41.608% |
| ecIspE | 39.051% | | 83.704% |
| kpIspE | 41.608% | 83.704% | |

**Figure S2**: **Sequence alignment of the IspE from *K. pneumoniae*, *E. coli*, *A. baumannii***. Secondary structure elements of IspE are indicated at top of the alignment; α helices and β strands are presented as curves and arrows. The identity, shown as a bar graph above the sequences, was calculated using the three aligned sequences. Amino acids in the active site are indicated by red boxes.

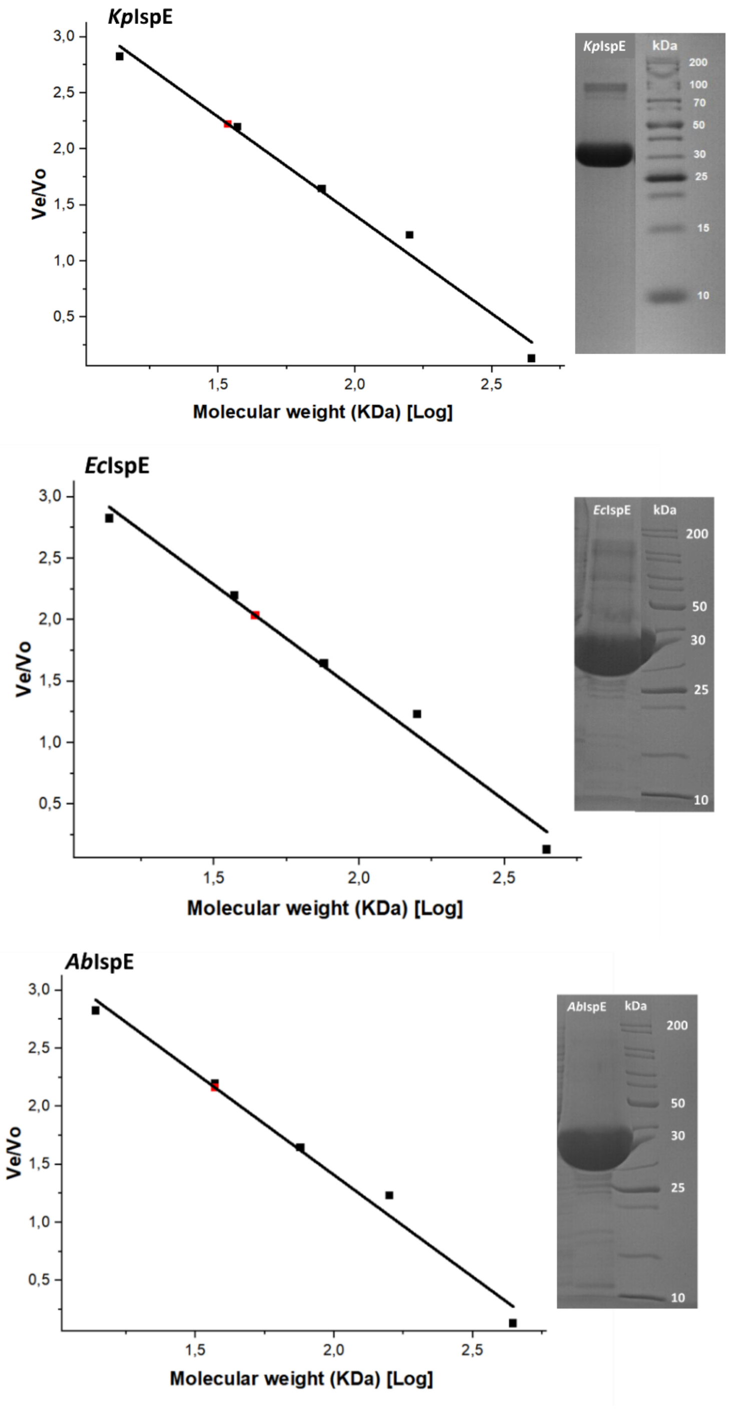


**Figure S3: A: Gel filtration calibration for molecular weights calculation and SDS-PAGE 12% electrophoresis** for *K. pneumoniae* IspE, *E. coli* IspE and *A. baumannii* IspE**.** Size exclusion chromatography column S200 16/600 was calibrated using Thyroglobulin (670 KDa), y-Globulin (158 KDa), ovalbumin (44 kDa), Myoglobin (17 KDa), and Vitamin B12 (1,350 KDa). The calibration curve was plotted using the elution volume/ column void volume (Ve/V0) versus logarithm of the molecular weight. Straight line is the calibration curve calculated from the data for molecular weight standards. Red dots correspond to the positions of Ve/Vo values for *K. pneumoniae IspE*, *E. coli IspE* and *A. baumannii IspE* respectively. Linear equation, from the calibration curve was used to calculate the experimental molecular weights of *K. pneumoniae IspE* (31 kDa), *E. coli IspE* (34 kDa) and *A. baumannii IspE* (30 kDa). SDS-PAGE 12% electrophoresis of IspEs showing the expected band size obtained purification.

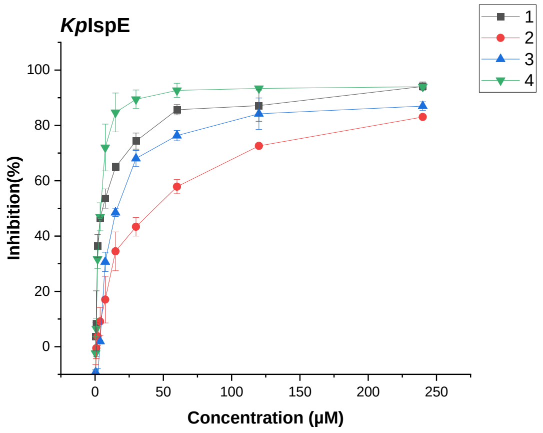


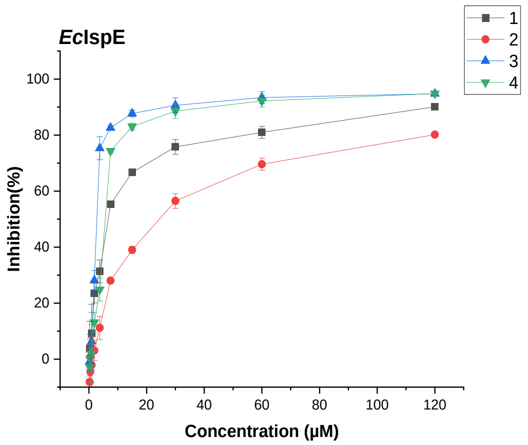


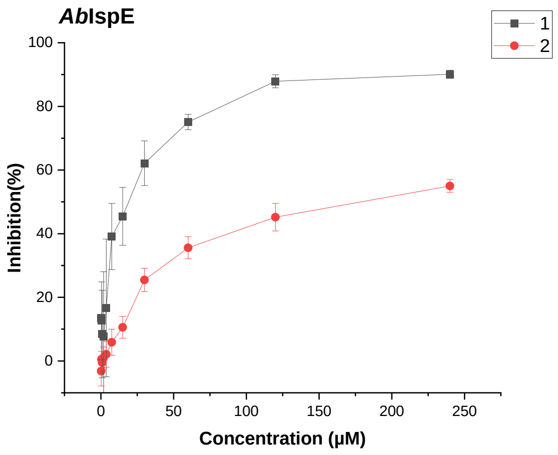


**Figure S4: Inhibitory dose-response curves** to determine the $IC_{50}$ for each inhibitor on *E. coli IspE (Ec*IspE*)*, *K. pneumoniae* IspE (*Kp*IspE) and *A. baumannii* IspE (*Ab*IspE). The curves were plotted using % of inhibition values of the enzymes after incubation with varied concentration of compounds. *E. coli* IspE concentration (200nM), *K. pneumoniae IspE* (150nM) and *A. baumannii* IspE (100nM). CDP-ME was used at (~1X $K_m$); 300 μM for *E. coli* IspE and *K. pneumoniae* IspE assay and 400 μM for *A. baumannii IspE*. ATP was kept at 1mM (~4X $K_m$) in all instances. The curves were fit using non-linear regression analysis program in OriginPro.

**Table S1:** Data collection and refinement statistics of the X-ray structures shown in the manuscript. Statistics for the highest-resolution shell are shown in parentheses

| | *K. pneumoniae* IspE | *E. coli* IspE (CDP-ME) | *E. coli* IspE (ligand 1) | *E. coli* IspE (ligand 2) | *E. coli* IspE (ligand 3) |
|---|---|---|---|---|---|
| PDB code | **8CKH** | **8QC7** | **8QCC** | **8QCN** | **8QCO** |
| Resolution range | 44.61 - 1.80 (1.968 - 1.80) | 35.75 - 2.03 (1.84 -2.03 ) | 47.93 - 1.904 (1.972 - 1.904) | 36.43 - 1.86 (1.922 - 1.856) | 48.28 - 1.55 (1.605 - 1.55) |
| Space group | P21 21 21 | C 1 2 1 | C 1 2 1 | C 1 2 1 | C 1 2 1 |
| Unit cell a b c [Å] α β γ [°] | 65.87 76.781 98.517 90 90 90 | 143.196 53.046 91.07 90 126.933 90 | 142.57 52.793 90.253 90 126.691 90 | 143.928 53.1129 91.7173 90 127.395 90 | 145.674 53.162 91.968 90 127.677 90 |
| Total reflections | 531648 (55829) | 207290 (20383) | 414388 (25077) | 510930 (35961) | 226873 (22755) |
| Unique reflections | 46367 (4526) | 35241 (3472) | 41807 (3718) | 45832 (4238) | 79873 (7952) |
| Multiplicity | 13.4 (14.2) | 4.4 (4.4) | 9.9 (6.7) | 11.1 (8.7) | 2.8 (2.9) |
| Wavelength | 0.97 | 0.93 | 0.93 | 0.93 | 0.62 |
| Completeness (%) | 98.9 (98.1) | 98.97 (98.52) | 98.41 (90.28) | 95.7 | 98.23 (98.82) |
| Mean I/ sigma (I) | 15.08 (4.28) | 5.21 (0.64) | 15.42 (1.01) | 3.61 (0.22) | 11.70 (0.99) |
| Wilson B-factor | 26.9 | 26.81 | 37.59 | 39.28 | 26.21 |
| R-merge | 0.07 (1.456) | 0.12 (2.184) | 0.078 (1.569) | 0.058 (1.341) | 0.046 (0.9965) |
| R-work | 0.1885 (0.2348) | 0.2165 (0.3117) | 0.1935 (0.3676) | 0.2271 (0.5133) | 0.2104 (0.3619) |
| R-free | 0.2267 (0.2870) | 0.2404 (0.3794) | 0.2353 (0.3785) | 0.2660 (0.5499) | 0.2411 (0.4087) |
| Number of non-hydrogen atoms | 4633 | 4627 | 4605 | 4523 | 4707 |
| Macromolecules | 4275 | 4334 | 4314 | 4330 | 4332 |
| Ligands | 88 | 192 | 165 | 138 | 150 |
| Solvent | 296 | 171 | 186 | 103 | 281 |
| Protein residues | 559 | 562 | 561 | 563 | 563 |
| RMS (bonds) | 0.005 | 0.020 | 0.041 | 0.024 | 0.031 |
| RMS (angles) | 0.80 | 0.75 | 1.27 | 0.66 | 1.17 |
| Average B-factor Protein | 33.65 | 33.24 | 41.86 | 48.57 | 33.14 |
| Solvent | 47.1 | 41.4 | 40.3 | 48.37 | 42.74 |
| Ligand (ADP) | 57.0 | 47.0 | 49.0 | 51.0 | 45.0 |
| Ligand (compound) | - | 35.0 | 37.0 | 49.0 | 37.0 |
| Ramachandran plot analysis (%) | | | | | |
| Favoured | 98.38 | 99.1 | 98.56 | 97.85 | 99.11 |
| Allowed | 1.62 | 0.90 | 1.44 | 2.15 | 0.89 |
| Outliers | 0.00 | 0.00 | 0.00 | 0.00 | 0.00 |

**Compounds synthesis:**

All reagents used for chemical synthesis were purchased from commercial suppliers, and used without further purification. All chemical yields refer to purified compounds and were not optimized. Reaction progress was monitored using TLC silica gel 60 $F_{254}$ aluminum sheets, and visualization was accomplished by UV at 254 nm. Column chromatography was performed using the automated flash chromatography system CombiFlash® Rf (Teledyne Isco) equipped with RediSepRf silica columns. Preparative RP-HPLC was was perofmed using an UltiMate 3000 Semi-Preparative System (Thermo Fisher Scientific) with nucleodur® C18 Gravity (250 mm × 10 mm, 5 µm) column. Purifications *via* preparative RP-HPLC were carried out using two possible conditions: *Condition A*: gradient 5–100% CH3CN + 0.05% HCOOH in water + 0.05% HCOOH in 53 min at a flow rate of 5 mL/min. The sample was dissolved in DMSO and manually injected to the HPLC system. *Condition B*: gradient 20–100% CH3CN + 0.05% HCOOH in water + 0.05% HCOOH in 35 min at a flow rate of 5 mL/min. Samples were dissolved in DMSO and manually injected to the HPLC system. $^1$H and $^{13}$C NMR spectra were recorded as indicated on a Bruker Avance Neo 500 MHz ($^1$H, 500 MHz; $^{13}$C, 126 MHz) with prodigy cryoprobe system or a Bruker Fourier 300 ($^1$H, 300 MHz; 13C, 75 MHz) instrument. Chemical shifts were recorded as δ values in ppm units and referenced against the residual solvent peak ($CDCl_3$, δ = 7.26, 77.16; DMSO-$d_6$, δ = 2.50, 39.52, MeOH-$d_4$: δ = 3.35, 4.78, 49.3). Splitting patterns describe apparent multiplicities and are designated as s (singlet), br s (broad singlet), d (doublet), dd (doublet of doublet), t (triplet), q (quartet), m (multiplet). Coupling constants (*J*) are given in hertz (Hz). Low resolution mass analytics and purity control of final compounds was carried out either using an Ultimate 3000-MSQ LCMS system (Thermo Fisher Scientific) consisting of a pump, an autosampler, MWD detector and a ESI quadrupole mass spectrometer. Purity of all compounds used in biological assays was ≥ 95%. High resolution mass spectra were recorded on a ThermoFisher Scientific (TF, Dreieich, Germany) Q Exactive Focus system equipped with heated electrospray ionization (HESI)-II source. Final products were dried at high vacuum.

**General Procedure A for the formation of a sulfonamide**:

The sulfonyl chloride (1 eq.) was added dropwise to a solution of propargyl amine (1 eq.) and Et3N (1.1 eq.) in dry CH2Cl2, at 0 °C. The mixture was left to stir at 25 °C for 30 min and concentrated in vacuo. Products were purified via column chromatography.

**General procedure B for the Sonogashira cross coupling**:

To an Ar-charged flask charged with iodocytosine (1.0 eq.), the previously synthesized alkyne (2 eq.), and Et3N (3.0 eq.) in anhydrous DMF, [PdCl2(PPh3)2] (0.1 eq.) and CuI (0.2 eq.) were added at 25 °C. The mixture was left to stir at 25 °C for 12 h. Water was added to the flask and the reaction mixture filtered, dissolved in DMSO and filtered again through celite. The reaction mixture was purified via HPLC.

**Chemical and Analytical Methods.**

$^1$H- and $^{13}$C-NMR spectra were recorded as indicated on a Bruker Avance Neo 500 MHz with prodigy cryoprobe system or a Bruker Fourier 300 instrument. Chemical shifts are given in parts per million (ppm), and referenced against the residual solvent peak. Coupling constants (J) are given in Hertz. Low-resolution mass analytics and purity control of final compounds were carried out either using an Ultimate 3000-MSQ LCMS system (Thermo Fisher Scientific), consisting of a pump, an autosampler, MWD detector and an ESI quadrupole mass spectrometer. High resolution mass spectra were recorded on a ThermoFisher Scientific (TF, Dreieich, Germany) Q Exactive Focus system equipped with heated electrospray ionization (HESI)-II source. Reagents were used as obtained from commercial suppliers without further purification. Procedures were not optimized regarding yield. Column chromatography was performed using the automated flash chromatography system

CombiFlash® Rf (Teledyne Isco) equipped with RediSepRf silica columns. Preparative HPLC was performed using an UltiMate 3000 Semi-Preparative System (Thermo Fisher Scientific) with nucleodur® C18 Gravity (250 mm × 10 mm, 5 µm) column. Final products were dried at high vacuum.

Derivatives of **1** were synthesized in three steps (Scheme S 1). First, sulfonamides were synthesized from the corresponding sulfonyl chloride and propargylamine. Iodination of cytosine was performed using $I_2$ and iodic acid to yield 5-iodocytosine. Finally, a sonogashira cross coupling reaction is performed, resulting in the target derivative.

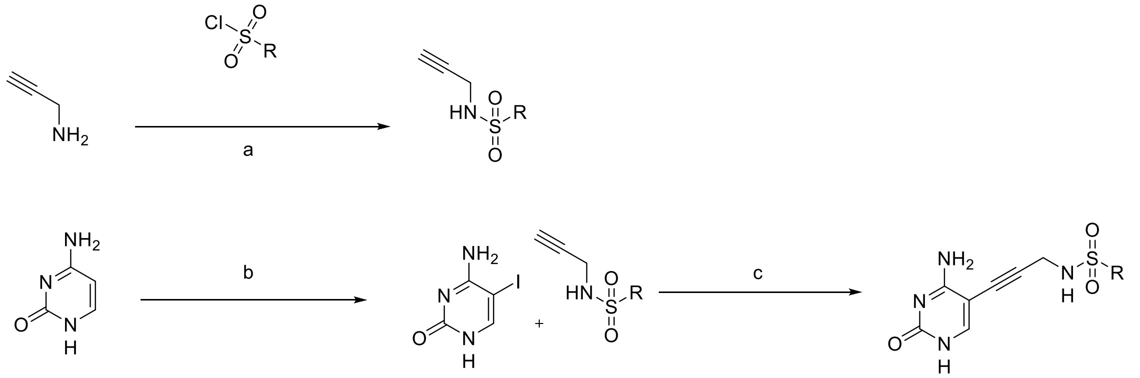


*Scheme S 1: Reagents and condition: a) TEA, DCM, 0°C-r.t, 30min; b) $I_2$, $HIO_3$, AcOH, 40°C, 12h; c) [PdCl2(PPh3)2], CuI, TEA, DMF, r.t, 4-12 h;*

**Synthesis of N-(3-(4-amino-2-oxo-1,2-dihydropyrimidin-5-yl)prop-2-yn-1-yl)cyclopropanesulfonamide (2)**

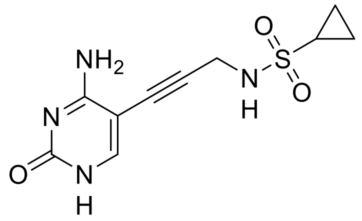

The title compound was synthesized according to the general method B. The crude product was purified by preparative HPLC (Condition B). The title product was obtained as a white solid, (70 mg, 0.26 mmol, 38%). $^1$H NMR (500 MHz, (DMSO-d$_6$) δ 8.52 (s, 1H), 4.16 (s, 2H), 1.56 (m, 1H), 0.69 (m, 4H). $^{13}$C NMR (126 MHz, (DMSO-d$_6$) δ 165.87, 155.54, 146.56, 142.20, 91.79, 76.12, 33.45, 30.07, 5.49. Purity: 100%; MS (ESI+) *m/z* 287 [*M*+H]+; HRMS (ESI+) *m/z* calcd for $C_{10}H_{12}N_4O_3S$ [*M*+H]+: 287.0630 found 287.0424.

**Synthesis of N-(3-(4-amino-2-oxo-1,2-dihydropyrimidin-5-yl)propyl)cyclopropanesulfonamide**

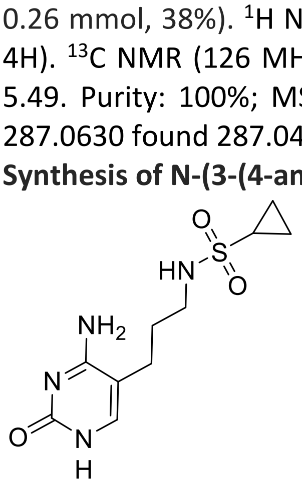

To a stirring solution of 1 (100 mg, 0.38 mmol, 1 eq.) in EtOH, Pd/C (10 mg, 0.1 mmol, 0.25 eq.) was added. $H_2$ gas was bubbled in to the stirring solution and the reaction mixture was allowed to stir for 3 h. The mixture was filtered over celite and washed with EtOH and concentrated under reduced pressure. The title product was obtained as a white solid, (92 mg, 0.34 mmol, 89%). $^{1}$H NMR (500 MHz, (DMSO-$d_6$) δ 7.24 (s, 1H), 3.05 (t, *J*=6.1 Hz, 2H), 3.05 (t, *J*=6.1 Hz, 2H), 2.59-2.57 (m, 1H), 2.33 (t, *J*=7.2 Hz, 2H), 1.66 (tt, *J*=7.4 Hz, 2H), 1.00-0.94 (m, 4H). $^{13}$C NMR (126 MHz, (DMSO-$d_6$) δ 157.39, 146.65, 142.28, 141.63, 44.47, 33.39, 27.99, 26.25, 13.71. Purity: 95.51%; MS (ESI+) *m/z 273* [*M*+H]+; HRMS (ESI+) *m/z* calcd for $C_{10}H_{16}N_4O_3S$ [*M*+H]+: 273.0943 found 273.1013.

**Synthesis of N-(3-(4-amino-2-oxo-1,2-dihydropyrimidin-5-yl)prop-2-yn-1-yl)cyclopentanesulfonamide (3)**

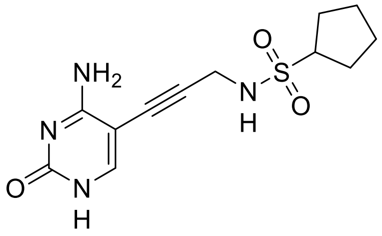

The title compound was synthesized according to the general method B. The crude product was purified by preparative HPLC (Condition B). The title product was obtained as a white solid, ( 74 mg, 0.25 mmol, 36%). $^{1}$H NMR (500 MHz, (DMSO-$d_6$) δ 7.67 (s, 1H), 4.03 (d, *J*= 5.6 HZ, 2H), 3.68-3.64 (m, 1H), 1.93-1.84 (m, 2H), 1.68-1.53 (m, 2H), 1.28-1.18 (m, 4H). $^{13}$C NMR (126 MHz, (DMSO-$d_6$) δ 167.59, 160.12, 140.16, 134.79, 87.08, 80.27, 54.41, 53.71, 33.34, 23.00. Purity: 100%; MS (ESI+) *m/z* 297 [*M*+H]+; HRMS (ESI+) *m/z* calcd for $C_{12}H_{16}N_4O_3S$ [*M*+H]+: 297.0943 found 297.1021.

**Synthesis of N-(3-(4-amino-2-oxo-1,2-dihydropyrimidin-5-yl)prop-2-yn-1-yl)cyclohexanesulfonamide (4)**

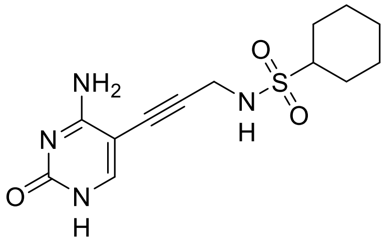

The title compound was synthesized according to the general method B. The crude product was purified by preparative HPLC (Condition B). The title product was obtained as a white solid, (8 mg, 0.026 mmol, 28%). $^{1}$H NMR (500 MHz, (DMSO-$d_6$) δ 7.67 (s, 1H), 4.00 (s, 2H), 3.07 (t, *J*= 8.7 Hz, 1H), 2.07-2.06 (m, 1H), 1.78-175 (m, 2H), 1.62-1.59 (m, 1H), 1.37-1.09 (m, 6H). $^{13}$C NMR (126 MHz, (DMSO-$d_6$) δ 159.03, 153.85, 141.32, 128.22, 48.49, 47.03, 31.41, 26.80, 23.90. Purity: 99.8%; MS (ESI+) *m/z* 311 [*M*+H]+; HRMS (ESI+) *m/z* calcd for $C_{13}H_{18}N_4O_3S$ [*M*+H]+: 311.1100 found 311.1770.

**Synthesis of N-(3-(4-amino-2-oxopyrimidin-1(2H)-yl)propyl)cyclopropanesulfonamide (HIPS7996)**

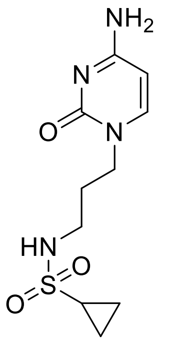

A solution of TEA (0.1 mL, 0.81 mmol. 1.2 eq) and of 4-amino-1-(3-aminopropyl) pyrimidin-2(1H)-one (115 mg, 0.68 mmol, 1 eq.) in DCM was cooled to 0 °C. Cyclopropanesulfonyl chloride (0.54 mL, 0.54 mmol, 0.8 eq.) was added to the solution with stirring. The reaction was allowed to warm to room temperature and stir for an addition 2 hours. The reaction mixture was concentrated and purified via HPLC (condition A). The title product was obtained as a white solid (14 mg, 0.05 mmol, 8%). $^1$H NMR (500 MHz, (DMSO-$d_6$) δ 7.98 (d, *J*= 7.6 Hz, 1H), 6.03 (d, *J*= 7.4 Hz, 1H), 3.82 (t, *J*= 6.8 Hz, 2H), 3.03 (q, *J*= 6.2, 6.5 Hz, 2H), 1.85-1.82 (m, 1H), 1.19 (t, *J*= 7.3 Hz, 2H), 0.95-0.89 (m, 2H), 0.57-0.50 (m, 2H). $^{13}$C NMR (126 MHz, (DMSO-$d_6$) δ 173.27, 162.36, 161.59, 125.01, 98.26, 96.90, 40.92, 29.53, 28.75, 10.20. Purity: 100%; MS (ESI+) *m/z* 272, [*M*+H]+; HRMS (ESI+) *m/z* calcd for $C_{10}H_{16}N_4O_3S$ [*M*+H]+: 273.3230 found 273.2943.

**Synthesis of N-(3-(4-amino-2-oxo-1,2-dihydropyrimidin-5-yl)prop-2-yn-1-yl)cyclopropanecarboxamide (HIPS7876)**

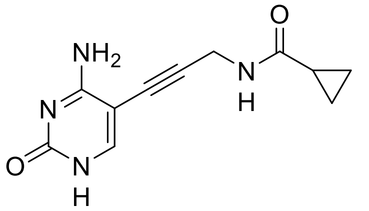

The title compound was synthesized according to the general method B. The crude product was purified by preparative HPLC (Condition A). The title product was obtained as a white solid, (73 mg, 0.3 mmol, 23%). $^1$H NMR (500 MHz, (DMSO-$d_6$) δ 7.67 (s, 1H), 4.04 (s, 2H), 2.66-2.61 (m, 1H), 0.97-0.95 (m, 4H). $^{13}$C NMR (126 MHz, (DMSO-$d_6$) δ 177.14, 155.40, 150.95, 136.22, 129.64, 85.37, 82.71, 32.81, 15.69, 9.37. Purity via HPLC: 100%; MS (ESI+) *m/z* 233 [*M*+H]+; HRMS (ESI+) *m/z* calcd for $C_{11}H_{12}N_4O_2$ [*M*+H]+: 233.0960 found 233.1031.

**N-(3-(4-amino-2-oxo-1-(tetrahydrothiophen-2-yl)-1,2-dihydropyrimidin-5-yl)prop-2-yn-1-yl)cyclopropanecarboxamide (HIPS8061)**

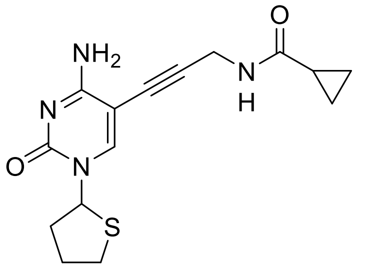

The title compound was synthesized according to the general method B. The crude product was purified by preparative HPLC (Condition A). The title productwas obtained as a white solid. (20 mg, 0.06 mmol, 10%). $^1$H NMR (500 MHz, (DMSO-$d_6$) δ 8.18 (s, 1H), 6.22 (dd, *J*= 5.6, 5.8 Hz, 1H), 4.18 (d, *J*= 5.0 Hz, 2H), 3.32 (ddd, *J*= 6.4, 6.0, 4.1 Hz, 1H), 2.91 (ddd, *J*= 6.4, 6.4, 3.7 Hz, 1H), 2.28-2.23 (m, 1H), 2.13-2.01 (m, 3H), 1.63 (tt, *J*= 5.3, 5.4 Hz, 1H), 0.75-0.73 (m, 4 H). $^{13}$C NMR (126 MHz, (DMSO-$d_6$) δ 172.47, 163.52, 161.33, 130.52, 126.88, 88.54, 83.51, 64.23, 44.07, 32.57, 28.65, 23.78, 15.84, 6.72. Purity via HPLC: 100%; MS (ESI+) *m/z* 319 [*M*+H]+; HRMS (ESI+) *m/z* calcd for $C_{15}H_{18}N_4O_2S$ [*M*+H]+: 319.1150 found 319.1218.

**Synthesis of N-(3-(4-aminopyrimidin-5-yl)prop-2-yn-1-yl)cyclopropanesulfonamide (HIPS8013)**

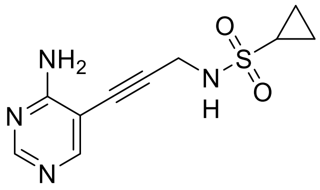

The title compound was synthesized according to the general method B. The crude product was purified by preparative HPLC (Condition B). The title product was obtained as a white solid, (5 mg, 0.02 mmol, 21%).$^{1}$H NMR (500 MHz, (DMSO-d$_6$) δ8.08 (d, *J*= 7.2 Hz, 1H), 7.14 (d, *J*= 7.3 Hz, 1H), 3.80 (t, *J*= 7.1 Hz, 2H), 3.09 (q, *J*= 6.8, 6.1 Hz, 2H), 1.78 (t, *J*= 6.9 Hz, 2H), 1.51-1.49 (m, 1H), 0.87-0.84 (m, 2H), 0.67-0.63 (m, 2H). $^{13}$C NMR (126 MHz, (DMSO-d$_6$) δ 154.91, 150.40, 129.30, 93.72, 47.34, 46.13, 29.50, 28.93, 5.08, 4.72. Purity: 100%; MS (ESI+) *m/z* 252, [*M*+H]+; HRMS (ESI+) *m/z* calcd for $C_{10}H_{12}N_4O_2S$ [*M*+H]+: 253.2920 found 253.0736.

**Synthesis of 6-Amino-5-iodo-2(1H)-pyrimidinone (5-iodocytosine)**

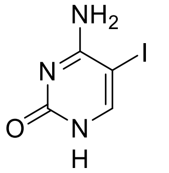

Cytosine (1 g, 9 mmol, 1 eq.), iodine (3.2 g, 13.5 mmol, 1.5 eq.) and iodic acid (2.2 g, 12.6 mmol, 1.4 eq.) were stirred in acetic acid at 40 °C overnight. The reaction mixture was cooled and treated with sat. aq. $Na_2S_2O_3$ until a white suspension was obtained. The mixture was neutralized with NaOH (10%). The product was collected via filtration, washed with cold water (5x) and concentrated under reduced pressure. _ was obtained as a white solid. Yield 97%. $^{1}$H NMR (500 MHz, ($CDCl_3$) δ 7.98 (s, 1H). MS (ESI+) *m/z* 237.

**Synthesis of 5-iodopyrimidin-4-amine**

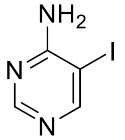

4-Aminopyrimidine (0.5 g, 5.25 mmol, 1 eq.), N-iodosuccinimide (1.2 mL, 12.6 mmol, 2.4 eq.) were stirred in acetic acid at 80 °C with stirring for 2.5 hours. The reaction mixture was cooled and extracted with DCM (3x). The organic layer was washed with sat. aq. $Na_2S_2O_3$ (3x) and brine (3x) and then concentrated under reduced pressure. The title product was obtained as a yellow solid. Yield 91%. $^{1}$H NMR (500 MHz, ($CDCl_3$) δ 8.44 (s, 1H), 8.30 (s, 1H).

**4-amino-5-iodo-1-(tetrahydrothiophen-2-yl)pyrimidin-2(1H)-one**

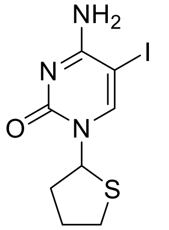

To a suspension of Synthesis of 5-iodopyrimidin-4-amine (0.30 g, 1.26 mmol, 1.0 eq) in a mixture toluene/DMF (2:1, 12 mL), triethylamine (0.70 mL, 5.06 mmol, 4 eq ) and TMSOTf (1.8 mL, 10.08 mmol, 8eq) were added. The mixture was stirred at rt for 15 min. Then, tetrahydrothiophene 1-oxide

(0.11 mL, 1.26 mmol, 1 eq) was added dropwise followed by additional triethylamine (1.05 mL, 7.56 mmol, 6eq). The mixture was stirred at rt for 30 min. The reaction was quenched by the addition of ice and extracted with EtOAc, followed by being washed with sat. aq. $NaHCO_3$ (3 x 10 mL). The crude product was purified by preparative HPLC (Condition B). The title compound was obtained as a white solid. Yield: 12 %. $^1$H NMR (500 MHz, (DMSO-$d_6$) δ 8.15 (s, 1H), 6.12 (dd, *J*= 6.4 Hz, 1H), 3.29-3.22 (m, 1H), 2.87-2.83 (m, 1H), 2.20-2.02 (m, 1H), 2.06-1.96 (m, 3H).

**Synthesis of N-(prop-2-yn-1-yl)cyclopropanesulfonamide**

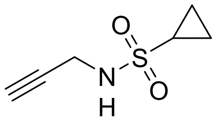


The title compound was synthesized according to the general method A. The crude product was purified by column chromatography using a gradient of 0-100% EtOAc in Hexanes to yield compounds _. Yield 69%. $^1$H NMR (500 MHz, ($CDCl_3$) δ 3.91 (dd, *J*= 2.5, 3.7 Hz, 2H), 2.51-2.46 (m, 1H), 2.28 (t, *J*= 2.4 Hz, 1H), 1.18-1.15 (m, 2H), 1.00-0.96 (m, 2H). MS (ESI+) *m/z* 160.

**Synthesis of N-(prop-2-yn-1-yl)cyclobutanesulfonamide**

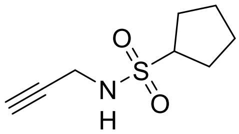


The title compound was synthesized according to the general method A. The crude product was used without any further purification.

**Synthesis of N-(prop-2-yn-1-yl)cyclohexanesulfonamide**

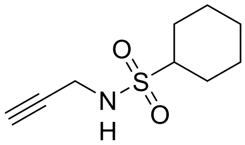


The title compound was synthesized according to the general method A. The crude product was purified by column chromatography using a gradient of 0-100% EtOAc in Hexanes to yield compounds _. Yield 35%; $^1$H NMR (500 MHz, ($CDCl_3$) δ 3.88 (dd, *J*= 2.5, 3.7 Hz, 2H), 2.97-2.92 (m, 1H), 2.27 (s, 1H), 2.17-2.14 (m, 2H), 1.86-1.83 (m, 2H), 1.66-1.63 (m, 1H), 1.49-1.44 (m, 5H), 1.26-1.12 (m, 2H); MS (ESI+) *m/z* 202.

**Synthesis of tert-butyl (3-bromopropyl)carbamate**

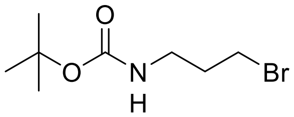


A mixture of 3-bromopropylamin hydrobromide (2 g, 9.14 mmol, 1 eq.), $(BOC)_2O$ (2.1 g, 9.59 mmol, 1.05 eq.) and triethyl amine (6.5 ml , 29.3 mmol,3.2 eq.) was stirred in dry DCM at room temperature for 24 h. The crude mixture was filtered and washed with diethyl ether to yield the product as a white solid. Yield 89%. $^1$H NMR (500 MHz, ($CDCl_3$) δ 3.56 (t, *J*= 6.7 Hz, 2H), 3.09 (t, *J*= 6.4 Hz, 2H), 1.97 (q, *J*= 6.5, 6.7 Hz, 2H), 1.44 (s, 9H).

**Synthesis of 4-amino-1-(3-aminopropyl)pyrimidin-2(1H)-one**

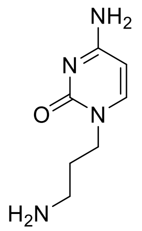

Cytosine (388 mg, 3.5 mmol, 1.2 eq.) and 60% NaH (147mg, 3.7 mmol, 1.05 eq.) were dissolved in dry DMF and stirred for 10 minutes. BOC-3-bromopropylamine (1 g, 4.2 mmol, 1 eq.) was added slowly and the reaction mixture was stirred at room temperature for 4 h. Water was added and the mixture was extracted with DCM (3x). The organic layer was concentrated in vacuum to obtain a yellow oil. The crude product was then dissolved in a methanol and acidified with HCl. The reaction mixture was stirred at room temperature for 1.5 hours. The reaction was concentrated under vaccum and dissolved in DCM. The crude mixture was washed with water (3x), brine (3x) and then purified via column chromatography. Yield 46%. $^{1}$H NMR (500 MHz, (DMSO-$d_6$) δ 7.50 (d, *J*= 7.0, 1H), 5.56 (d, *J*= 7.1, 1H), 3.53 (t, *J*= 6.9 Hz, 2H), 3.03 (q, *J*= 6.6, 6.6.1 Hz, 2H), 1.58 (t, *J*= 6.7 Hz, 2H).

**NMR Spectra:**

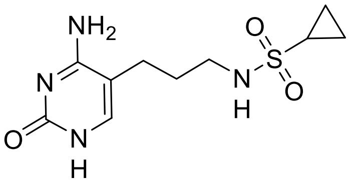

$H^1$ NMR spectrum:

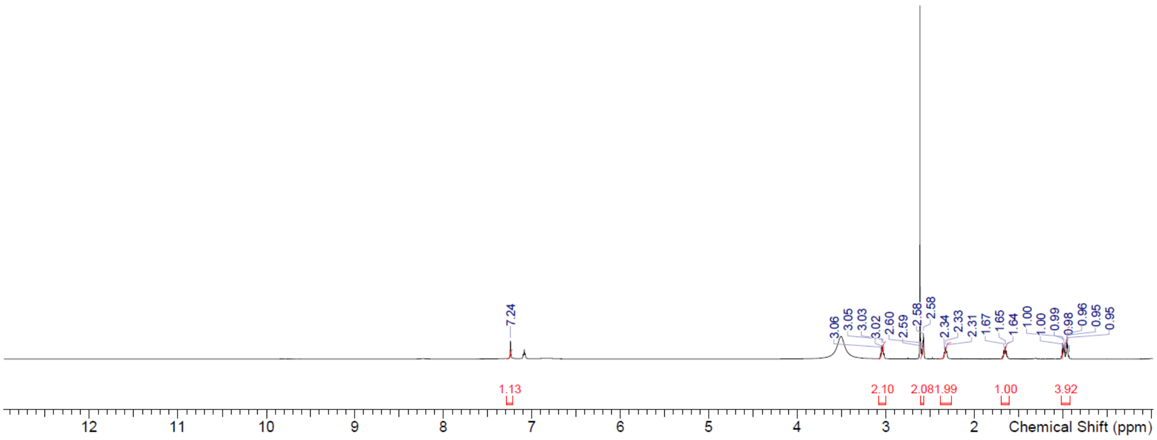


$C^{13}$ NMR spectrum:

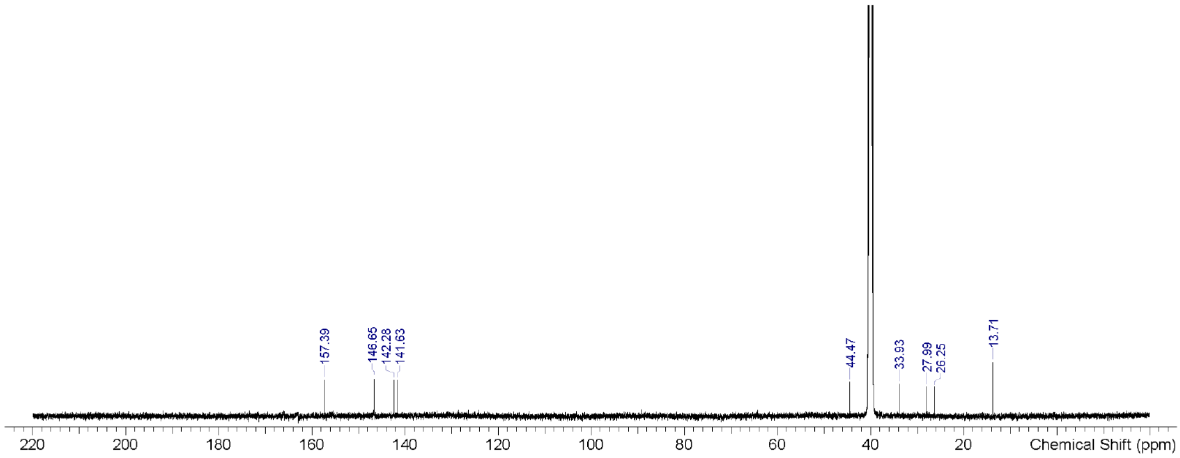

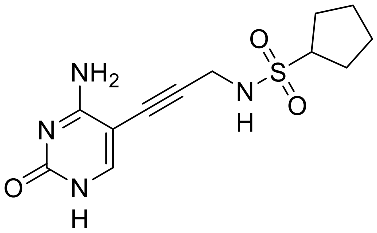


H[1] NMR spectrum:

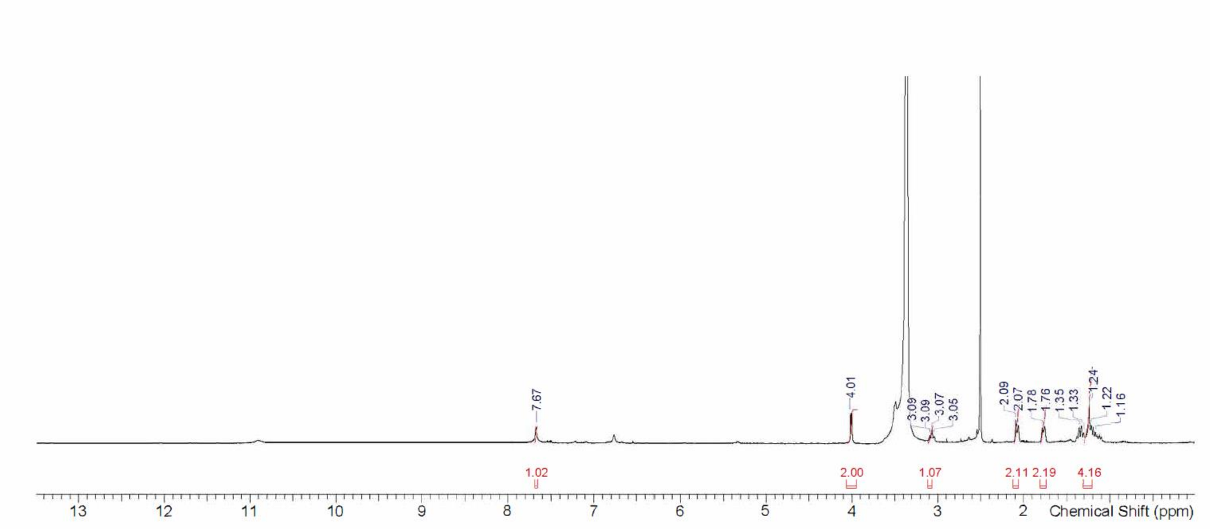


C[13] NMR spectrum:

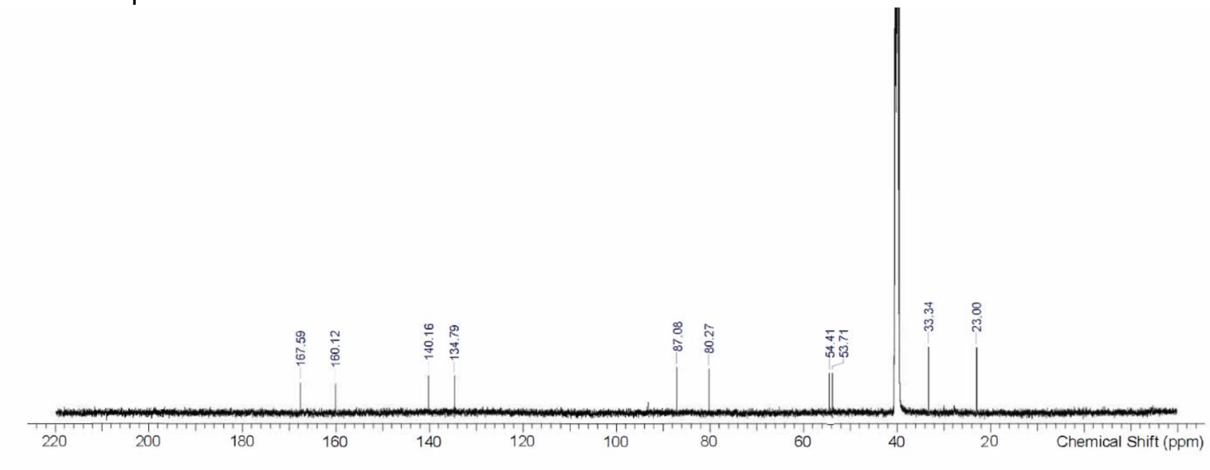

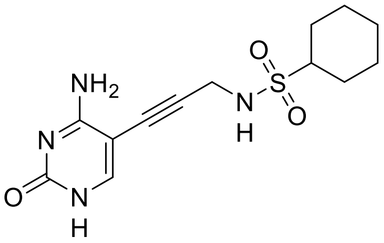


$H^1$ NMR spectrum:

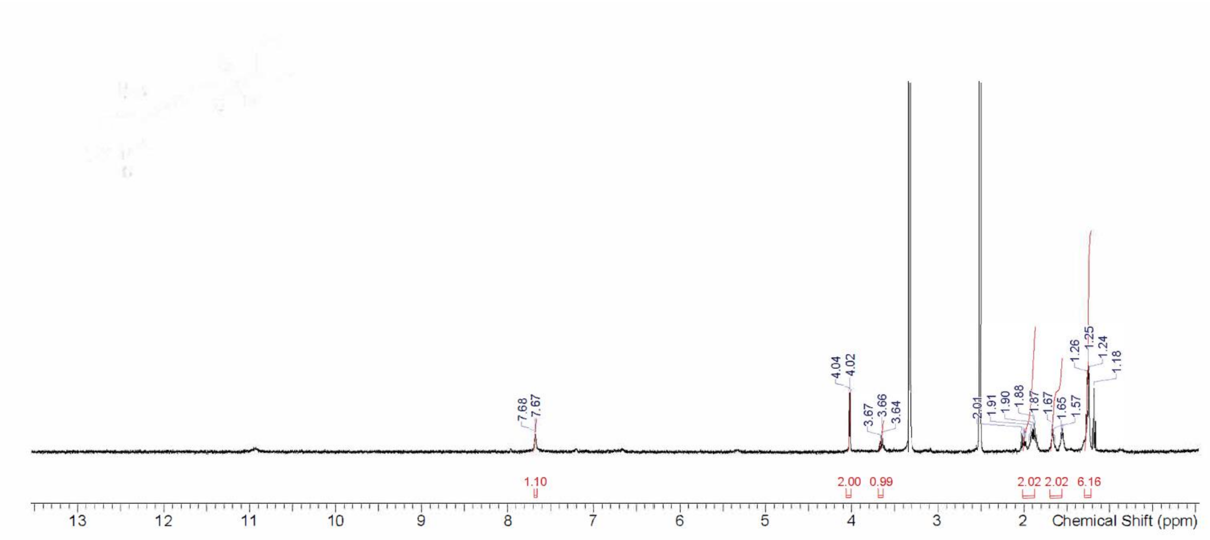


$C^{13}$ NMR spectrum:

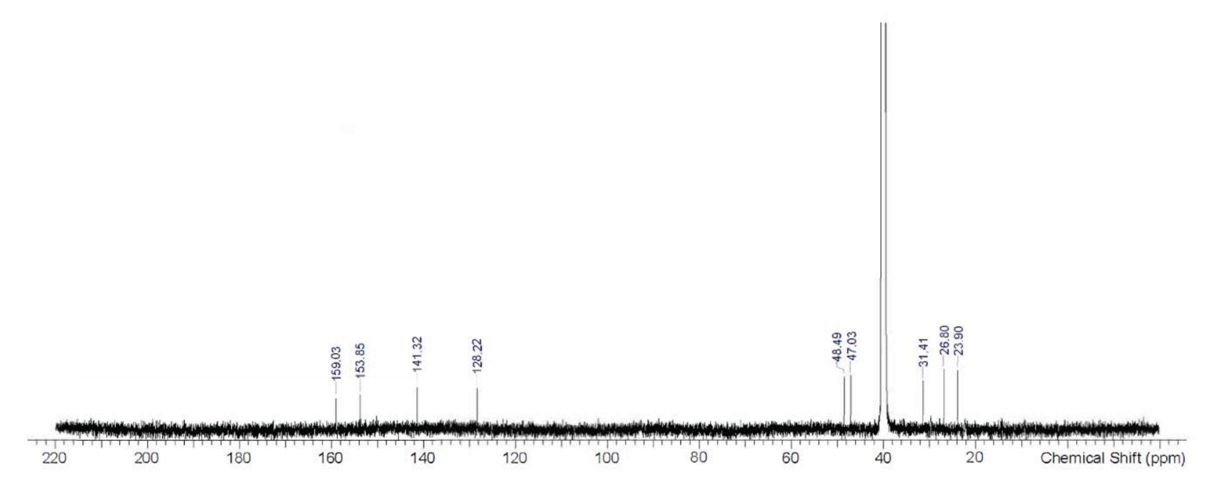

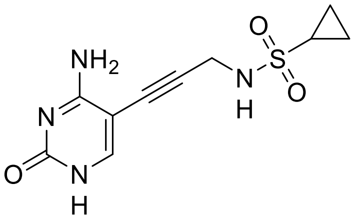


$H^1$ NMR spectrum:

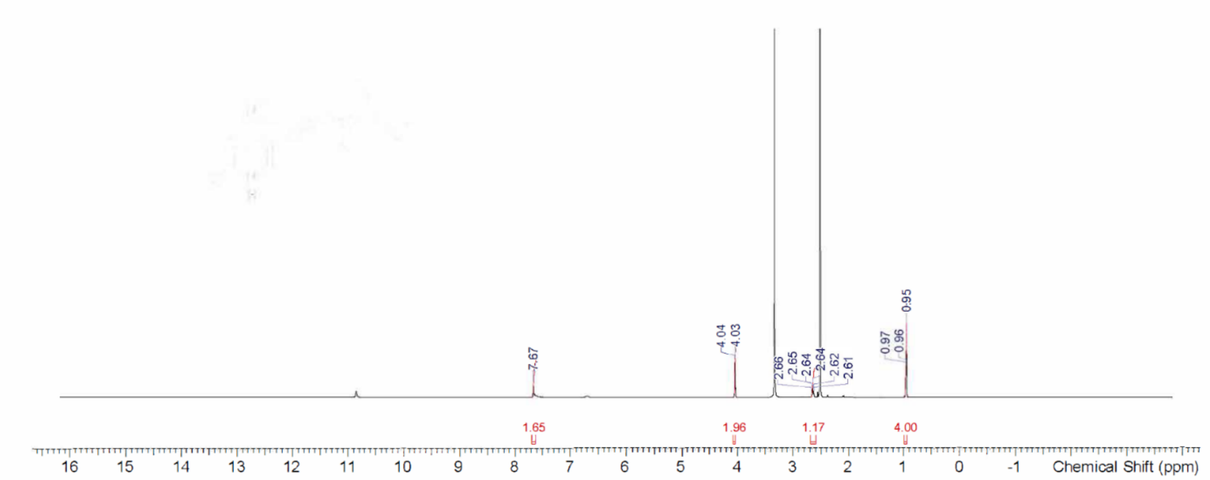


$C^{13}$ NMR spectrum:

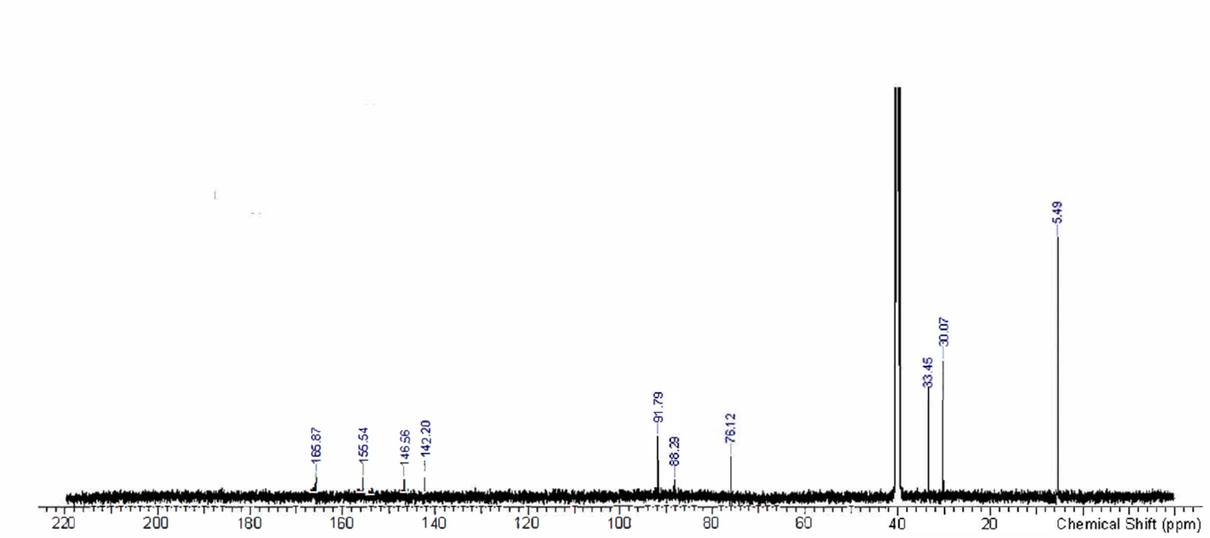

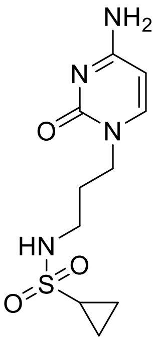


H1 NMR spectum:

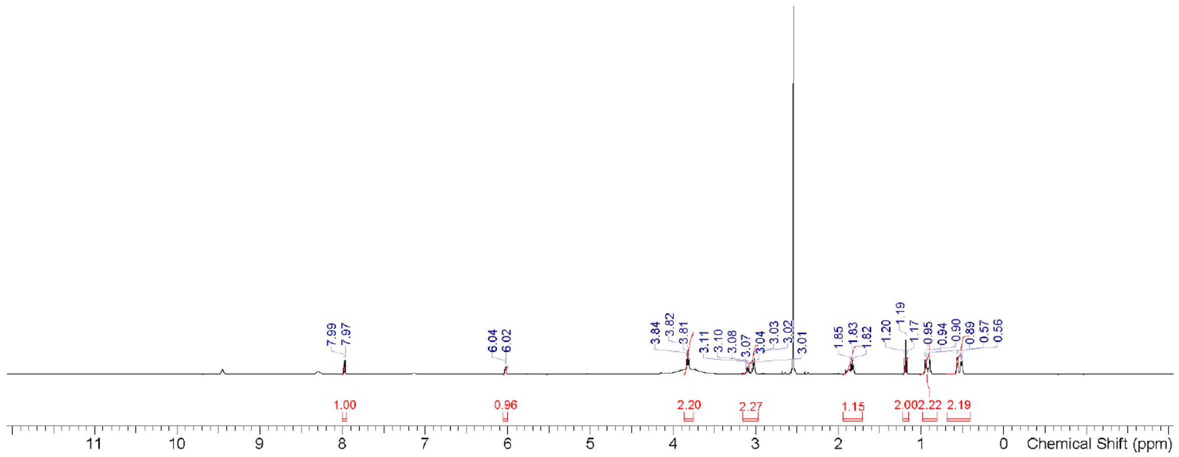


C13 NMR spectrum:

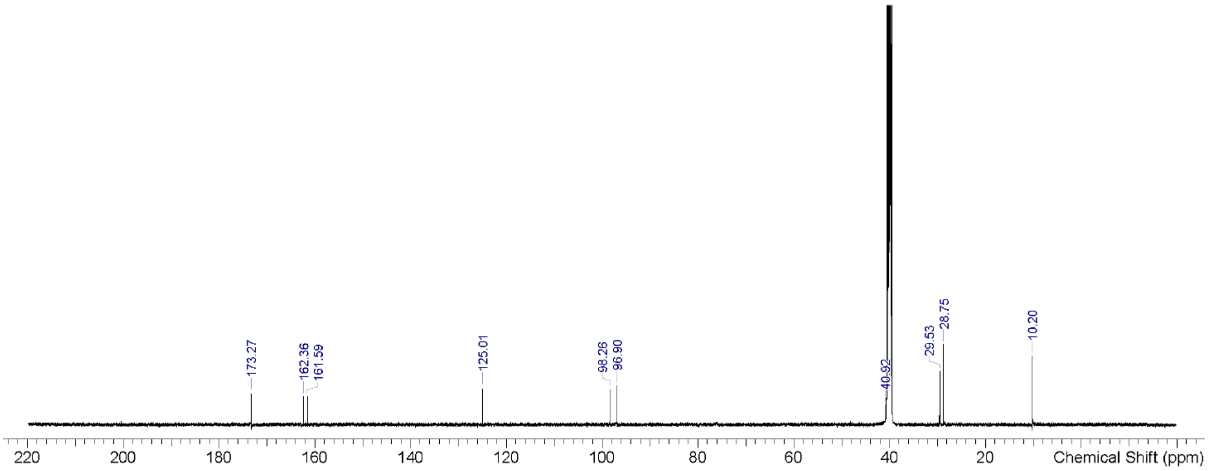

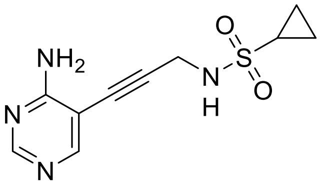


H1 NMR spectrum:

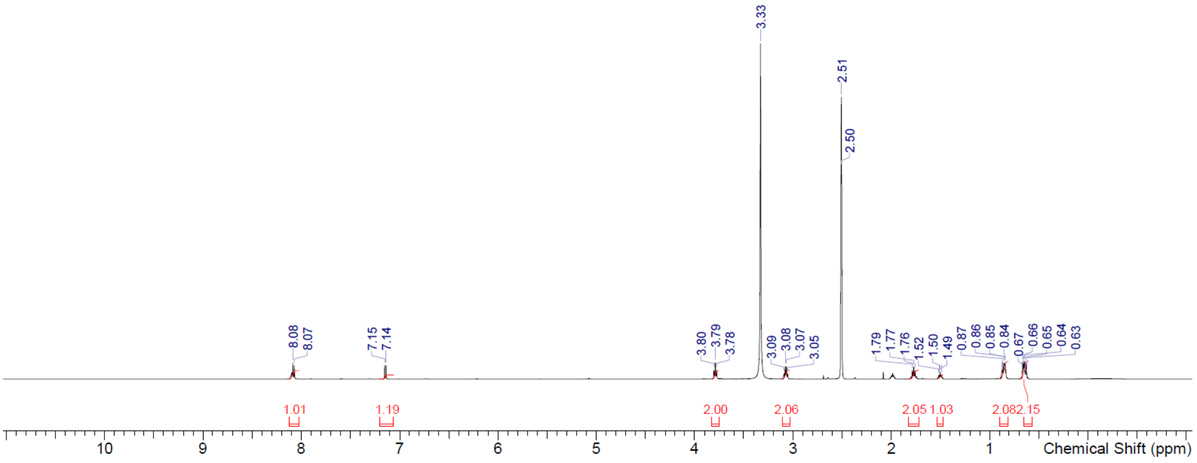


C13 NMR spectrum:

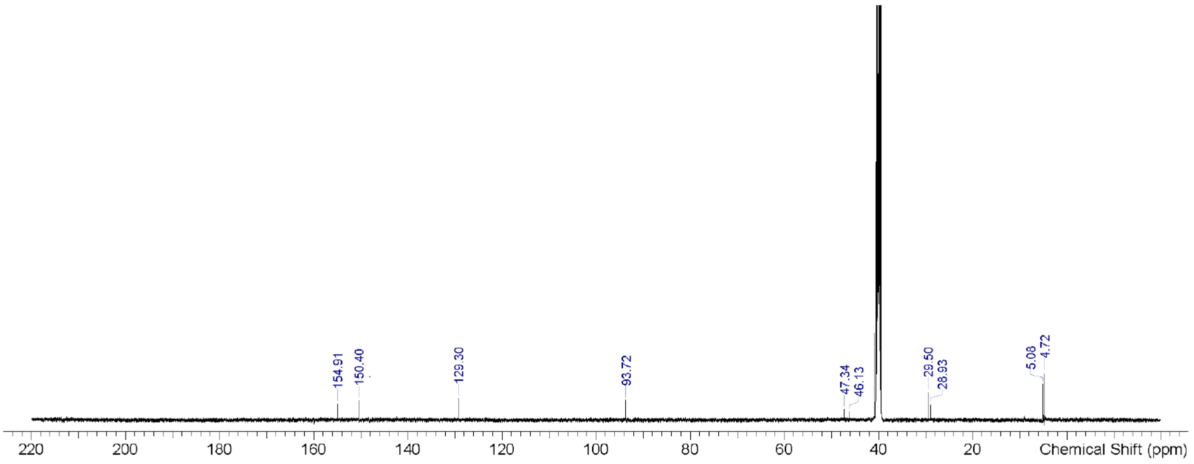

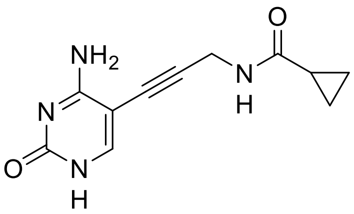


H1 NMR

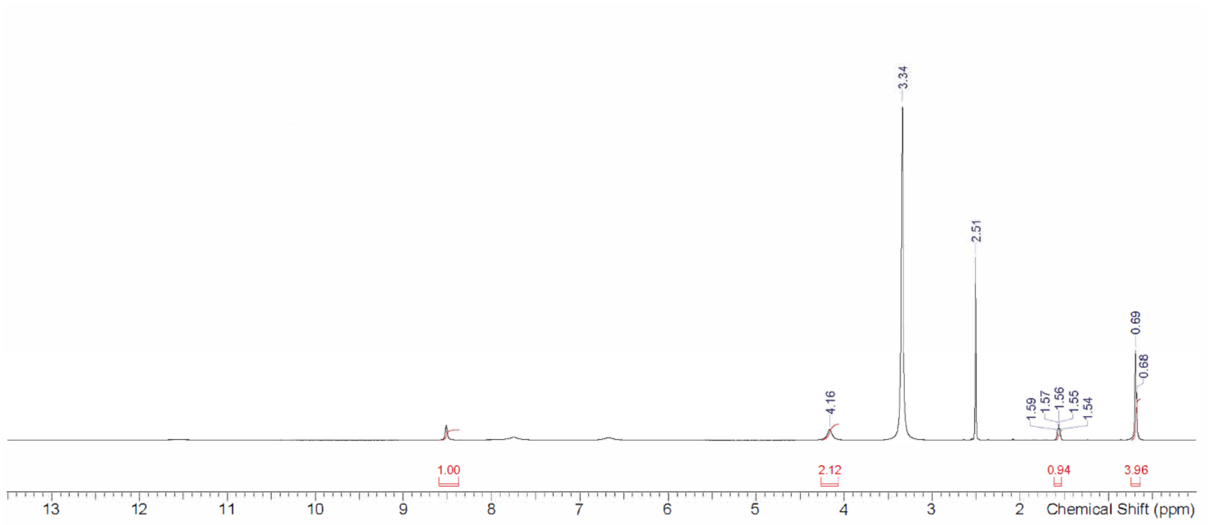


C13 NMR

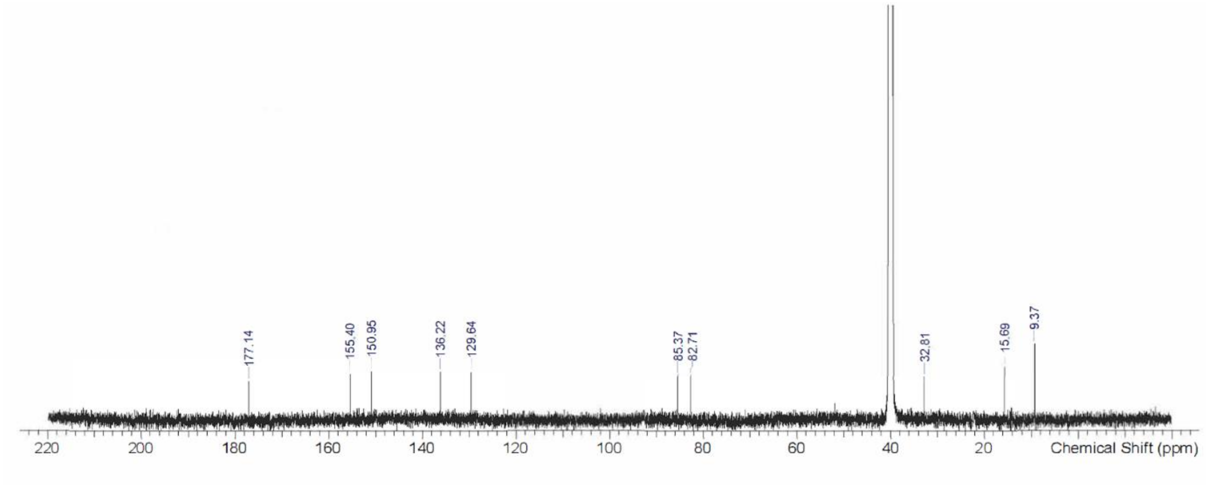

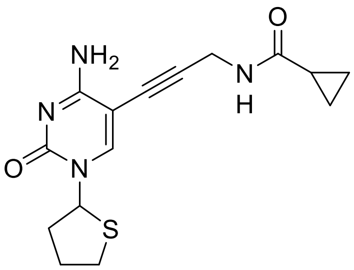


H1 NMR

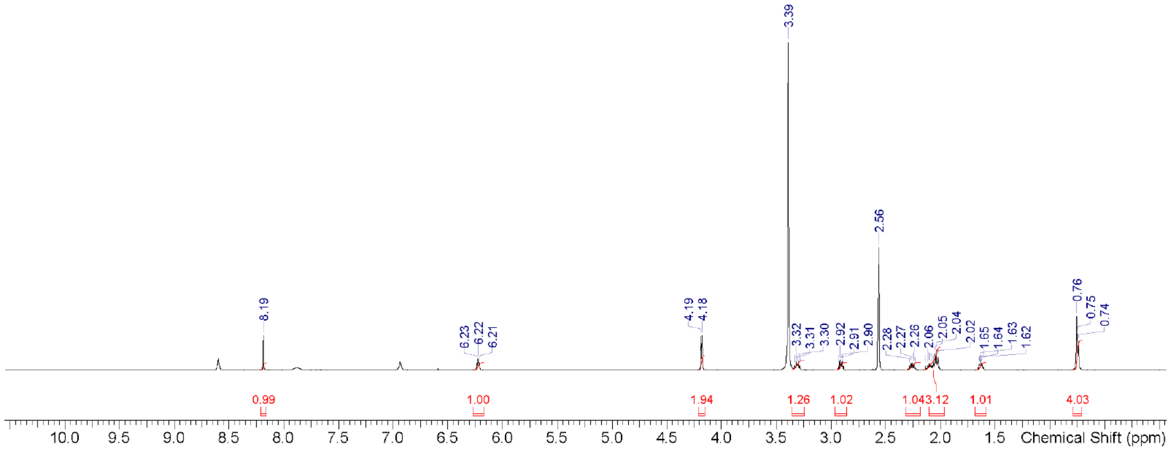


C13 NMR

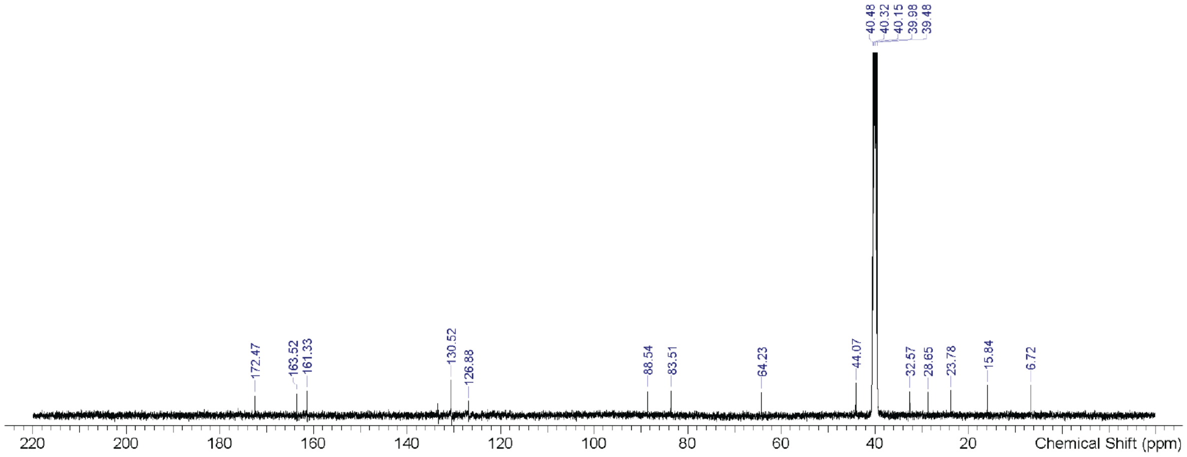

**Computational methods: Docking of IspE inhibitors to *E. coli IspE* and *A. baumannii IspE***

All subsequent steps were carried out in MOE v2020.09.

**Preparation of ligands.** The 2D structures of the three compounds (**2, 3 & 4**) were sketched using ChemDraw professional 20.0 and were imported into the MOE window. The compounds were subjected to an energy minimization up to a gradient of 0.001 kcal $mol^{-1}$ $Å^2$ using the MMFF94x force field and R-field solvation model, then they were saved as a .mdb file. The predominant protonation status of the compounds in aqueous medium at pH 7 was calculated via the compute | molecule | wash command in the database viewer window.

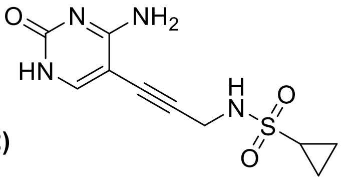

**(2)**

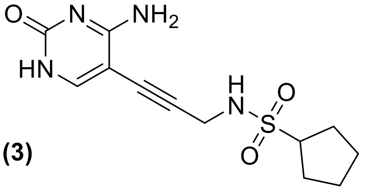

**(3)**

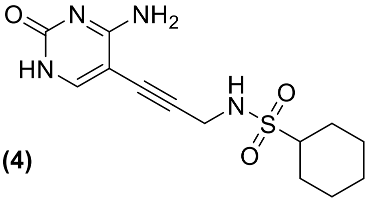

**(4)**

**Preparation of protein structures.**

<u>*E. coli* IspE</u>

The X-ray co-crystal structure of *E. coli IspE* with **3** was used for the molecular docking studies. Potential was set up to Amber14:EHT as a force field and R-field for solvation. Missing residues and termini were fixed using the Structure Preparation module in MOE. The Protonate3D module was used to assign and fix hydrogens atoms. Only the receptor atoms were selected in this case as this step would cause the loss of aromaticity in the cytosine ring of the co-crystalized ligand when the ligand was included in the Protonate3D calculation. Tethered energy minimization was then performed via the QuickPrep module, with the Structure Preparation and Protonate3D tick boxes unchecked (as these were already performed manually) while using the default settings for the rest of the module.

<u>*A. baumannii IspE* (Uniprot accession code : B2HUM5)</u>

The Swiss-Model structure was downloaded as a .pdb file from the Swiss-Model website via the link provided on the Uniprot web page. Potential was set up to Amber14:EHT as a force field and R-field for solvation. Correction of library errors, and tethered energy minimization of binding site were performed via the QuickPrep module using default settings. The structure was then aligned with the *E. coli IspE* structure in MOE using the sequence viewer's alignment module ("Sequence and Structural" sequence alignment and "Use Current Alignment" superposition methods) with an overall RMSD of 1.118 Å.

**Binding site selection.**

For *E. coli* IspE, the binding site was defined as the residues within 4.5 Å of the co-crystalized ligand. For the *A. baumannii* IspE structure, the equivalent residues (as determined by the sequence alignment) were selected as the binding site for docking.

**Molecular docking**

<u>*E. coli IspE*</u>

Molecular docking was performed via the Dock module of MOE. The co-crystalized ligand was selected as a template for docking (similarity mode) while the receptor was set to the prepared *E. coli IspE* structure and the site was selected as described above. Triangle matcher placement was used with 100 poses followed by Rigid Receptor Refinement with 10 poses. The GBVI/WSA dG scoring function was used for both methods with default settings.

<u>*A. baumannii IspE*</u>

Molecular docking was performed via the Dock module of MOE. The default docking mode was used while the receptor was set to the prepared *E. coli IspE* structure and the site was selected as described above. Triangle matcher placement was used with 100 poses followed by Rigid Receptor Refinement with 10 poses. The GBVI/WSA dG scoring function was used for both methods with default settings.

**Table S 2.** Docking scores of the three compounds docked to the three structures of *E. coli IspE* and *A. baumannii* IspE.

| Compound | *E. coli* IspE Docking Score (kcal/mol) | *A. baumannii IspE* Docking Score (kcal/mol) |
|---|---|---|
| 2 | -6.95 | -6.1 |
| 3 | -7.08 | -6.23 |
| 4 | -7.04 | -6.6 |
